# Dynamic structural order of a low complexity domain facilitates assembly of intermediate filaments

**DOI:** 10.1101/2020.04.01.013870

**Authors:** Vasily O. Sysoev, Masato Kato, Lillian Sutherland, Rong Hu, Steven L. McKnight, Dylan T. Murray

**Affiliations:** Department of Biochemistry, UT Southwestern Medical Center, Dallas, TX, 75390; Department of Chemistry, University of California, Davis, CA, 95616

## Abstract

The coiled-coil domains of intermediate filament (IF) proteins are flanked by regions of low sequence complexity. Whereas IF coiled-coil domains assume dimeric and tetrameric conformations on their own, maturation of eight tetramers into cylindrical IFs is dependent upon either “head” or “tail” domains of low sequence complexity. Here we confirm that the tail domain required for assembly of *Drosophila* Tm1 IFs functions by forming labile cross-β interactions. These interactions are seen in polymers made from the tail domain alone as well as assembled IFs formed by the intact Tm1 protein. The ability to visualize such interactions *in situ* within the context of a discrete cellular assembly lends support to the concept that equivalent interactions may be used in organizing other dynamic aspects of cell morphology.

**One Sentence Summary:** A new form of protein folding that interconverts between the structured and unstructured states controls assembly of intermediate filaments.

## Main Text

Deposition of germ granules at the posterior tip of *Drosophila melanogaster* oocytes specifies formation of cells of the germinal lineage (*1*). Forward genetic studies have illuminated the importance of a number of genes essential to germ cell formation. Many of these genes encode RNA binding proteins of which RNA granules are themselves composed (*2-5*). Perplexingly, mutations proximal to the locus encoding the fly tropomyosin gene also impede deposition of germ granules and the subsequent formation of germ cells (*6*). These mutations interfere with the formation of a non-muscle isoform of tropomyosin designated Tm1 that is somehow required for germ cell specification (*7, 8*).

A program of alternative pre-mRNA splicing allows fly oocytes to produce an isoform of tropomyosin that replaces the domains necessary for interaction with troponin, and resultant sensitivity to regulation of muscle contraction by free calcium, with protein segments of low sequence complexity. Previously reported experiments have shown that the oocyte-specific Tm1 isoform assembles into intermediate filaments, and that its C-terminal low complexity (LC) domain is essential for filament assembly (*7*). Here we present a combination of biochemical and biophysical studies giving evidence that the C-terminal LC domain of Tm1 facilitates assembly of intermediate filaments by forming structurally specific cross-β interactions that are unusually dynamic. These studies of the fly Tm1 isoform of tropomyosin may be relevant to the behavior of prototypic intermediate filaments long studied in a variety of more complex organisms. They may further be instructive as to the behavior of LC domains in many other aspects of cell morphology.

## Results

An oocyte-specific program of alternative pre-mRNA splicing allows production of an isoform of tropomyosin, designated Tm1, that is markedly different from that made in muscle cells. Whereas the two proteins share sequences specifying a central, coiled-coil domain, alternative splicing yields a unique C-terminal tail domain within the oocyte-specific Tm1 isoform. This tail domain is 69 residues in length, of low sequence complexity (18 asparagine residues, 15 serine residues and 6 threonine residues, Supplementary data, Fig. S1), evolutionarily limited to the *Sophophora* sub-genus of *Drosophila*, and required for the protein to assemble into intermediate filaments (*7*).

Expression, purification and incubation of the isolated Tm1 tail domain leads to the assembly of amyloid-like polymers. X-ray diffraction studies of Tm1 tail domain polymers have revealed a prominent diffraction ring at 4.7Å. These and other studies led to the conclusion that the Tm1 tail domain can, in an isolated state, assemble into amyloid-like polymers that are labile to disassembly (*7*). In this regard, the Tm1 tail domain displays properties similar to the low complexity (LC) domains of numerous RNA binding proteins (*9, 10*), the FG repeats forming the permeability filter of nucleopores (*11-14*), and six different intermediate filament proteins expressed in mammalian cells and tissues (*15*).

### The Tm1 tail domain adopts a similar structural conformation in isolation and in assembled intermediate filaments

Uniform labeling of the Tm1 tail domain with ^13^C and ^15^N isotopes allowed analysis of amyloid-like polymers by solid-state NMR (ss-NMR) spectroscopy. That the polymers adopt a partially ordered conformation is confirmed by the spectra shown in Fig. 1A. When analyzed by ss-NMR at 16°C, the Tm1 tail domain-only polymer sample gave rise to strong signals in the spectrum recorded using ^1^H-^13^C cross polarization (CP) and high power ^1^H decoupling (*16, 17*). The spectrum likewise gave intense signals when recorded using ^1^H-^13^C insensitive nuclei enhanced by polarization transfer (INEPT) without ^1^H decoupling (*18*). Signals appearing in the CP-based spectrum arise from immobilized regions of the protein in the polymers. Signals appearing in the INEPT-based spectrum arise from highly mobile regions of the protein in the polymers. These data indicate that there are both well-ordered and conformationally dynamic regions within Tm1 tail domain polymers.

**Fig. 1.**
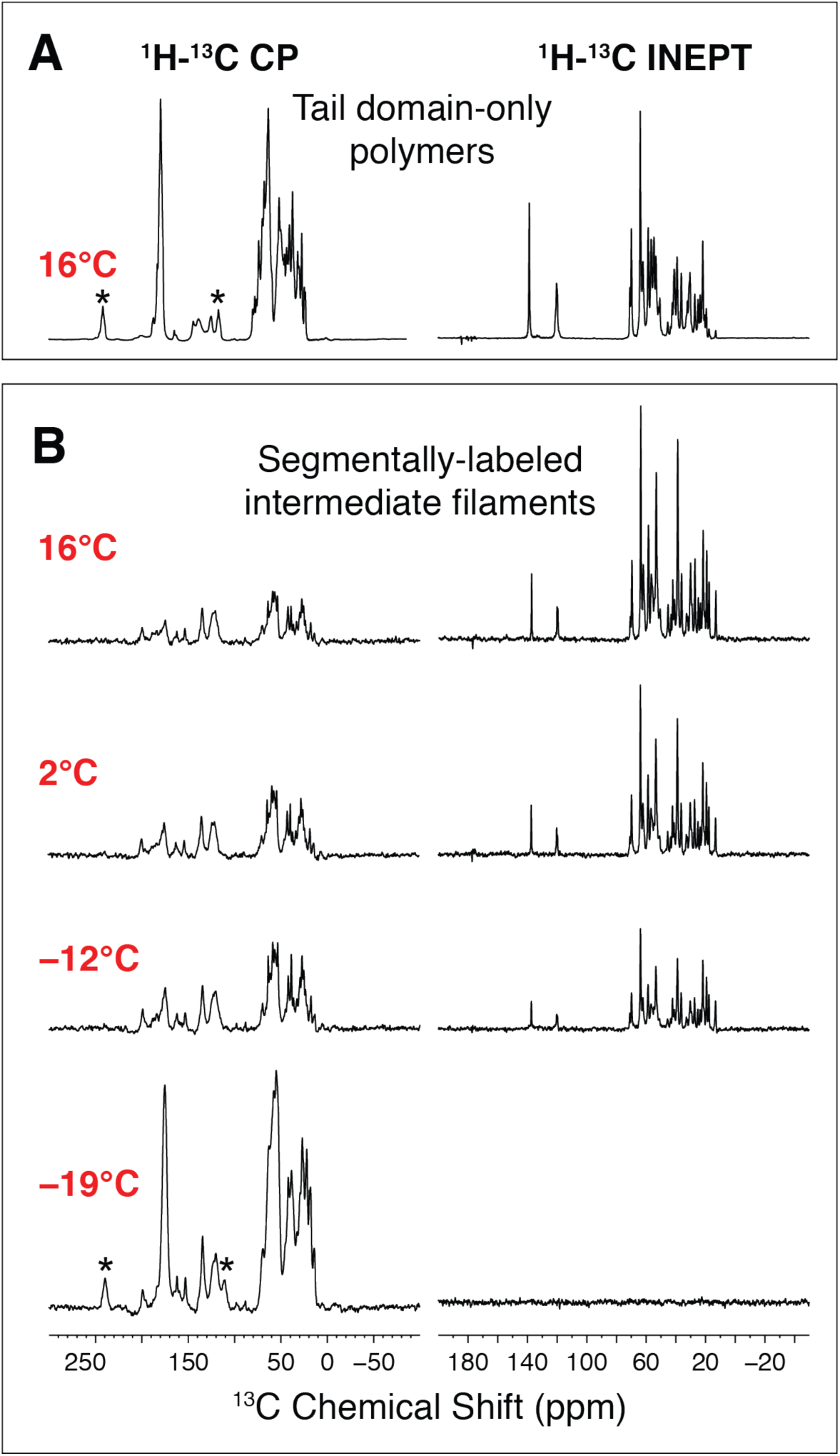
One-dimensional cross polarization- and INEPT-based solid state NMR spectra of Tm1 tail domain. Top panel (A) shows cross polarization (left) and INEPT (right) spectra of ^13^C/^15^N-labeled Tm1 tail domain-only polymers recorded at 16°C. Four sets of spectra shown below (B) represent cross polarization (left) and INEPT (right) spectra of segmentally ^13^C/^15^N-labeled Tm1 protein that was assembled into mature intermediate filaments. Isotopic labeling was restricted to the Tm1 tail domain by the use of intein chemistry as diagrammed in Supplementary Data, Fig. S2A. Cross polarization and INEPT spectra of fully assembled, segmentally labeled Tm1 intermediate filaments were recorded at 16, 2, −12 and −19°C as displayed top-to-bottom. Asterisks in the top and bottom cross polarization spectra indicate magic angle spinning side bands.

In order to study the conformation of the Tm1 tail domain in the context of fully assembled intermediate filaments, we used a split-intein system to produce a segmentally labeled form of the intact Tm1 protein (*19*). The tail domain, including residues 373 to 441, was expressed separately from the N-terminus of the protein (residues 1 to 372). The two segments were chemically ligated via a reaction scheme depicted in Supplementary Data, Fig. S2, yielding a full-length Tm1 protein containing a single asparagine-to-cysteine change at residue 371. This procedure allowed selective incorporation of ^13^C and ^15^N isotopes into the tail domain for the purpose of spectroscopically probing both its structure and dynamics within the full-length Tm1 protein as it exists in assembled intermediate filaments. The native Tm1 protein assembles into intermediate filaments having a diameter of 13 to 16 nm (*7*). Supplementary Data, Fig. S2 shows that segmentally, ^13^C/^15^N-labeled Tm1 intermediate filaments are morphologically indistinguishable from those assembled from the native protein as viewed by electron microscopy.

Assembled, segmentally ^13^C/^15^N-labeled Tm1 intermediate filaments were packed by centrifugation into an NMR rotor and analyzed at temperatures ranging from 16 to −19°C (Fig. 1B). Weak cross polarization (CP) signals for the sample analyzed at 16°C indicate that few regions within the Tm1 tail domain are sufficiently immobilized to allow significant CP magnetization transfers. The strong signals in the INEPT-based spectrum at 16°C are indicative of highly dynamic regions within the Tm1 tail domain as it exists in assembled intermediate filaments. As a function of temperature reduction to 2, −12 and −19°C, the strength of CP signals increased and that of INEPT signals decreased. These observations are consistent with a reduction in molecular motion of the Tm1 tail domain in intermediate filament assemblies as the sample temperature was reduced. The overall profile of the CP-based spectra of the Tm1 tail domain as viewed in isolated, tail domain-only polymers is similar to its conformation in ^13^C/^15^N-labeled intermediate filaments as assayed at low temperatures (Fig. 1). These 1D spectra are not, however, adequate to report on the actual molecular structure of the tail domain in the two different assemblies.

In order to more carefully probe both the uniformly ^13^C/^15^N-labeled Tm1 tail domain-only polymers and assembled, segmentally labeled Tm1 intermediate filaments, we performed two-dimensional ss-NMR to measure ^13^C chemical shifts from rigid sites within the assemblies. Fig. 2A shows a 2D ^13^C-^13^C CP-based dipolar assisted rotational resonance (CP-DARR) (*20, 21*) spectrum with 50 millisecond mixing time for the Tm1 tail domain-only polymers. Off-diagonal cross-peaks in CP-DARR spectra arise from rigid and structured carbon atoms that are one to three bonds apart in the protein. The spectrum of the Tm1 tail domain-only polymer sample exhibited sharp NMR resonances typical of a well-ordered protein structure with full-width-half-maximum (FWHM) linewidths of 175 to 225 Hz (0.8 to 1.2 ppm). Signals attributable to threonine, serine, isoleucine, cysteine, asparagine, aspartic acid, lysine, glutamic acid and alanine are clearly present in the spectrum. The ^13^C chemical shift values of these signals are consistent with β-strand conformations for all sites that could be unambiguously assigned to a single amino acid type.

**Fig. 2.**
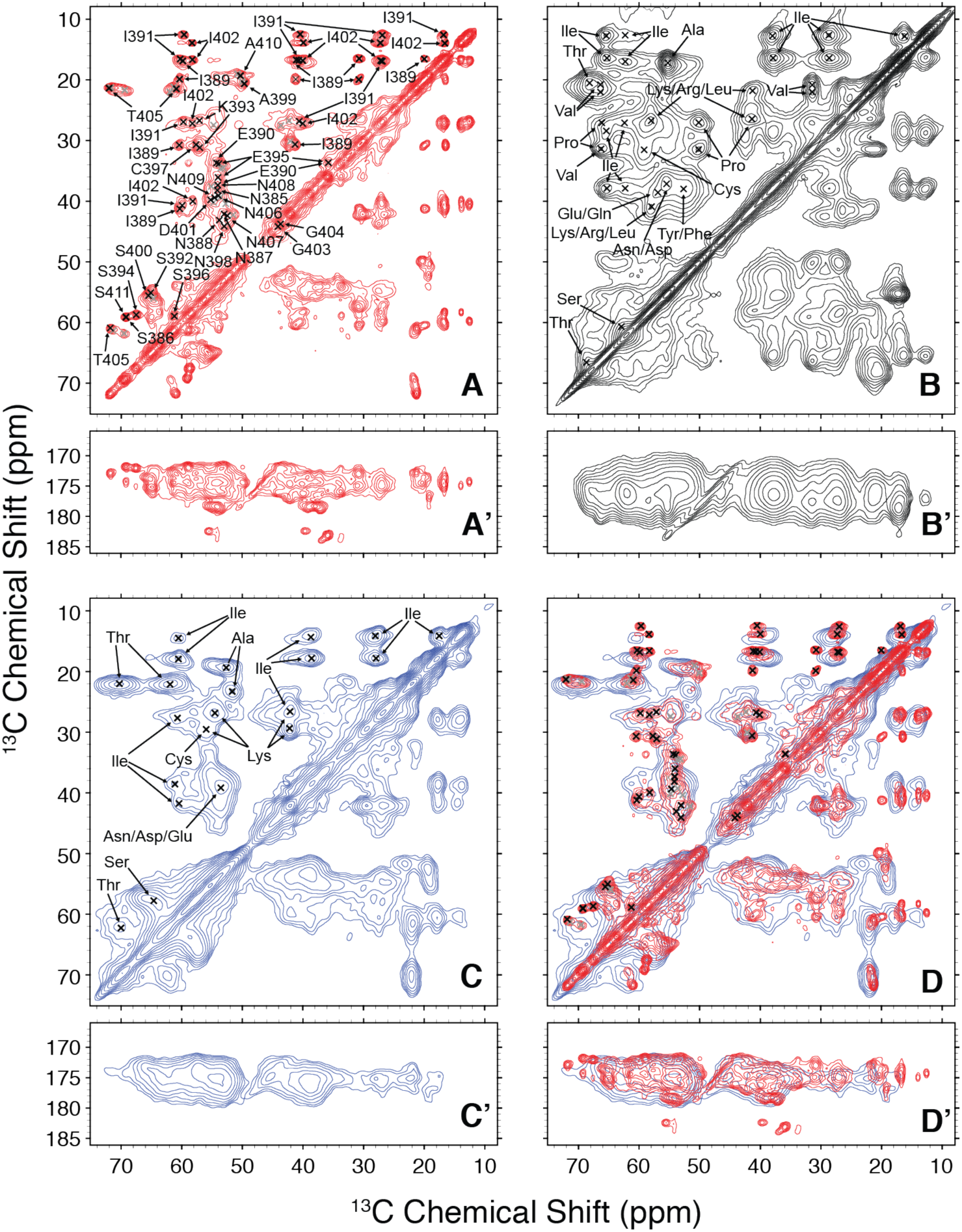
Two-dimensional cross polarization-based solid state NMR spectra of Tm1 tail domain. Top left panels (A and A’) show a ^13^C-^13^C CP-DARR spectrum with 50 millisecond mixing time of ^13^C/^15^N-labeled Tm1 tail domain-only polymers recorded at 16°C. Sequence-specific resonance assignments determined in this work are indicated by amino acid type and location within the full length Tm1 protein. Top right panels (B and B’) show the same spectrum of an ether precipitated sample of the Tm1 tail domain as analyzed in its polymeric form (panel A). Amino acid residues specified in the spectrum of ether precipitated protein were not assigned sequence-specifically. Bottom left panels (C and C’) show ^13^C-^13^C CP-DARR spectrum with 50 millisecond mixing time of segmentally ^13^C/^15^N-labeled Tm1 protein that had been assembled into intermediate filaments recorded at −19°C (see Supplementary Data, Fig. S2). Amino acid residues specified in the spectrum of segmentally labeled Tm1 intermediate filaments were not assigned sequence-specifically. Bottom right panels (D and D’) represent an overlay of the spectra shown in panels A/A’ and C/C’. The extensive overlap of signal intensity in the two samples gives evidence that the Tm1 tail domain exists in highly similar structural states in the two assemblies. Sequence-specifically assigned amino acids from the data shown in panels A/A’ are indicated by small X’s in the overlay of panels D/D’. The spectra of the Tm1 tail domain within assembled intermediate filaments (C and C’) overlap poorly or not at all with the spectra of ether-precipitated Tm1 tail domain (B and B’). An overlay of the latter two spectra is shown in Supplementary Data, Fig. S3.

That the chemical shift peaks observed in Tm1 tail domain polymers may be diagnostic of *bona fide* biologic structure is supported by ss-NMR spectra taken from an ether-precipitated sample of the same protein (Fig. 2B). Whereas the ether-precipitated sample revealed chemical shifts for numerous amino acid side chains within the Tm1 tail domain, few such peaks overlapped with those observed for tail domain-only polymers. The most noticeable differences in the respective spectra shown in Figs. 2A and 2B are significant reductions of serine and threonine β-strand signals, the appearance of a set of proline and valine signals, and a shift of the CA position of isoleucine signals.

The 2D ^13^C-^13^C CP-DARR spectrum of intermediate filaments prepared from the full-length, segmentally labeled sample of Tm1 exhibited broad NMR resonances with FWHM linewidths greater than 300 Hz at −19°C (Fig. 2C). It could not be determined from this spectrum alone if the intensity in the spectrum arose from: (i) multiple, single-site resonances with relatively narrow linewidths arising from multiple sites with well-defined conformations; or (ii) a single site with a broad line shape, indicating a significant degree of conformational heterogeneity. For either of these cases, the center positions of the signal intensities were consistent with β-strand conformations for the residues giving rise to the signals.

Despite the uncertainty specified above, the spectrum in Fig. 2C is consistent with a dynamic β-strand conformation for the Tm1 tail domain in the intermediate filament assemblies analyzed at 16°C, with slowed molecular motions as recorded at the reduced temperature of –19°C. An overlay of the 2D ^13^C-^13^C CP-DARR spectra of the isolated Tm1 tail domain-only polymers recorded at 16°C, and that of the tail domain as recorded at −19°C in segmentally labeled intermediate filaments, is shown in Fig. 2D. The overlay shows that the positions of CP signal intensities are highly similar between the two spectra. These similarities indicate that the Tm1 tail domain in the intermediate filament assemblies, when evaluated at reduced temperature, converges toward a structural conformation similar to that of the Tm1 tail domain-only polymers. Differences between the two spectra include one missing set of isoleucine peaks and slight differences in the ^13^C chemical shifts of the CG1 and CG2 cross-peaks for isoleucine residues. In stark contrast, an overlay of the spectra of the segmentally labeled Tm1 protein in its assembled state (Fig. 2C) and in the ether-precipitated state of the Tm1 tail domain (Fig. 2B) shows dramatic differences (Supplementary Data, Fig. S3). Despite being chemically identical, the Tm1 tail domain exhibits obvious structural differences in these two states.

### Structural characterization of the cross-β core-forming segment of the Tm1 tail domain

A set of 2D and 3D CP-based ^15^N-^13^C correlation spectra with 50 millisecond ^13^C-^13^C DARR mixing collected on the Tm1 tail domain-only polymer sample at 16°C allowed identification and structural characterization of the core forming segment of the polymers. Figs. 3A and 3A’ show the CP-based 2D NCACX spectrum of the Tm1 tail domain polymers (*22*). The black and grey X’s on the spectrum in Fig. 3A’ indicate 39 sets of signals observed in the 3D implementation of the NCACX experiment. Signals that have been sequence-specifically assigned are indicated by black X’s and labels in Fig. 3A’ (see Materials and Methods and Supplementary Data, Table S1).

**Fig. 3.**
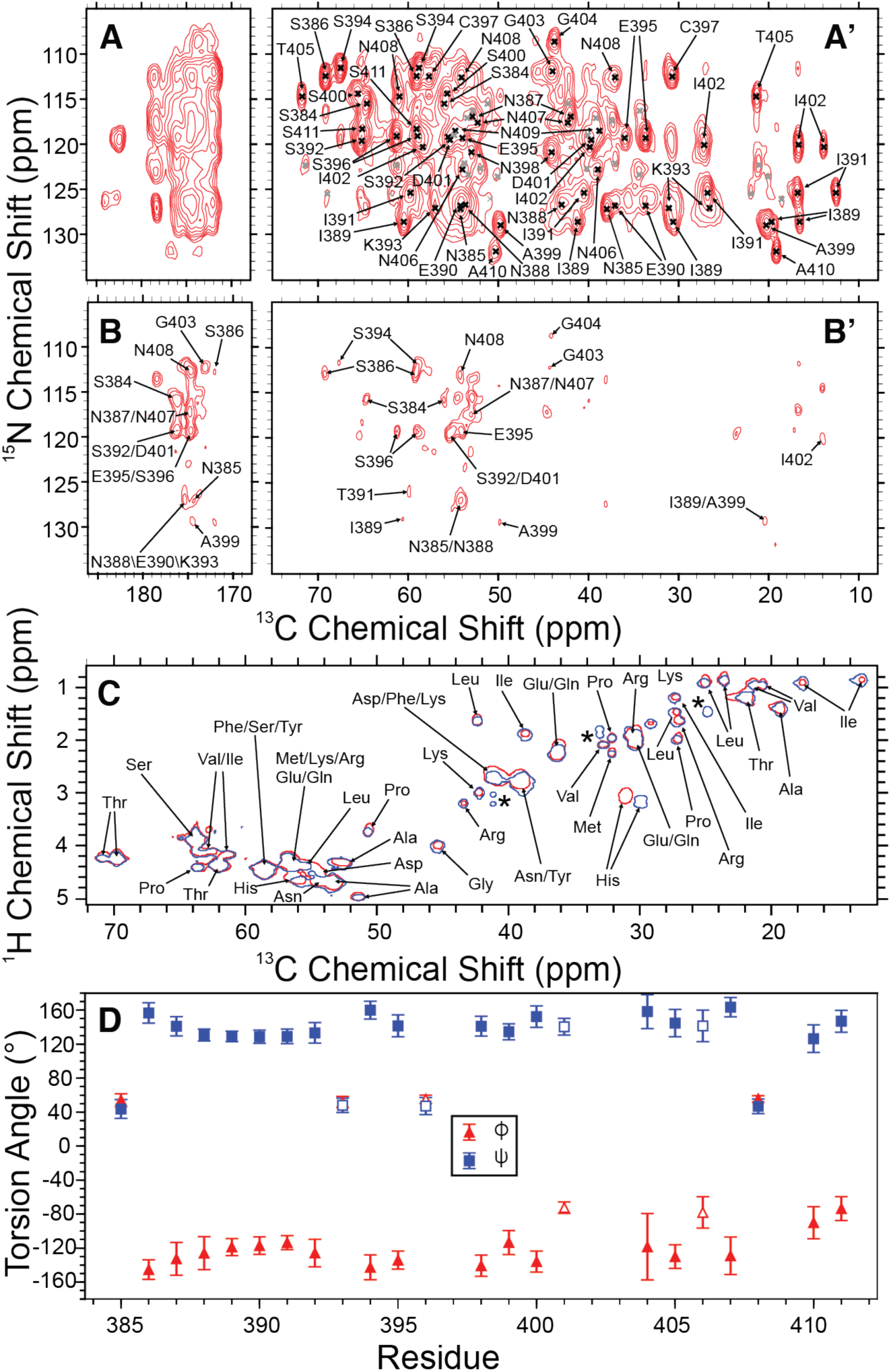
Structural characterization of Tm1 tail domain polymers. Top panels (A and A’) show a two-dimensional cross polarization-based NCACX spectrum of ^13^C/^15^N-labeled Tm1 tail domain-only polymers measured at 16°C. Panels immediately below (B and B’) show a two-dimensional ^15^N-^13^C zf-TEDOR spectrum of mixed (^13^C/^14^N)- and (^12^C/^15^N)-labeled Tm1 tail domain-only polymers measured at 5°C. Panel C shows two-dimensional ^1^H-^13^C INEPT-MAS spectra of ^13^C/^15^N-labeled Tm1 tail domain-only polymers (red contours) overlapped with segmentally ^13^C/^15^N-labeled Tm1 intermediate filaments (blue contours). Both spectra shown in panel C were recorded at 16°C. Bottom panel (D) shows NMR chemical shift based TALOS-N torsion angle predictions. Solid triangles and squares represent “strong” β-strand predictions and open triangles and squares represent “generous” β-strand predictions determined according to published criteria (*30*).

These assignments were obtained using four CP-based spectra: 2D and 3D NCACX, and 2D and 3D NCOCX (*22*). Representative 2D slices from the 3D NCACX spectrum are shown in Supplementary Data, Fig. S4. The 2D NCOCX spectrum and representative slices from the 3D NCOCX spectrum are shown in Supplementary Data, Fig. S5. Of the 39 sets of signals observed in these spectra, 38 can be unambiguously assigned to a single amino acid type. Complete lists of the observed NCACX signals are shown in Supplementary Tables S1 and S2. Two alanine, three isoleucine, eight asparagine, one lysine, two glutamic acid, seven serine, one threonine, one cysteine and two glycine residues were sequence-specifically assigned to residues 384 to 411 of the Tm1 tail domain. These amino acid residues represent roughly 30% of the total amino acid content of the Tm1 tail domain sequence.

Our assignment procedure was performed computationally using the MCASSIGN algorithm (*23*) in an iterative fashion comparable to previous work on FUS and hnRNPA2 LC domain polymers (*24, 25*) to yield statistically significant and unambiguous assignments for residues 384 to 411. A complete description of the assignment procedure is presented in Materials and Methods. Of note, no signals were consistently assigned to any region of the N-terminal His-tag sequence throughout the assignment calculations. The remaining unassigned signals have either relatively low signal-to-noise (values of ∼10 compared to the assigned signals which mostly range from ∼30 to ∼60) or correspond to strong signals from asparagine residues ambiguously assigned to the stretch of five consecutive asparagines located between positions 379 and 383. The low signal-to-noise peaks arise from alanine or threonine residues that are present in the unassigned region immediately following residue 411. Supplementary Data, Tables S1 and S2 list the assigned and unassigned signals with chemical shift and signal-to-noise values from the 3D NCACX spectrum.

A ^15^N-^13^C zf-TEDOR spectrum (*26*) of the Tm1 tail domain polymers at 5°C is shown in Fig. 3B. For this experiment, the Tm1 tail domain polymers were prepared from a solution of protein where 50% of the molecules were uniformly ^15^N isotopically labeled and 50% of the monomers were uniformly ^13^C isotopically labeled. The Tm1 tail domain polymers that result from this preparation should have, on average, alternating ^13^C- and ^15^N-labeled molecules. Signals in this spectrum arise from ^13^C and ^15^N nuclei that are 5Å or less apart in space (*26*). In the aliphatic carbon region, we observed signals with ^13^C and ^15^N chemical shifts consistent with the assignments for S384, N385, S386, N387, N388, I389, E390, T391, S392, S394, E395, A399, D401, I402, G403, G404, N407, and N408. In the carbonyl carbon region, we observed ^13^C and ^15^N chemical shifts consistent with the assignments for residues S384, N385, N387, N388, E390, S392, K393, E395, S396, A399, S400, D401, G403, T405, N407, and N408. The observed signal-to-noise ratios between 5 and 15 are consistent with the signals arising from long-range, 4 to 5Å ^13^C-^15^N distances during the 6.0 millisecond mixing time used in this experiment (*26, 27*). Of these signals, the only set of ^13^CA, ^13^CO, and amide ^15^N chemical shift values that were ambiguous between two different residues were for S392 and D401. The unambiguous signals observed in the spectrum in Fig. 3B are consistent with the same residues in adjacent monomers in the Tm1 polymers having ^15^N and ^13^C nuclei 4 to 5Å apart (*26*) (*i*.*e*. the ^13^C and ^15^N chemical shifts are assigned to the same amino acid). Slices illustrating the signal-to-noise in the zf-TEDOR spectrum are shown in Supplementary Data, Fig. S6.

Figure 3C shows an overlay of 2D INEPT-based ^1^H-^13^C spectra of the Tm1 tail domain at 16°C (red) and the assembled, segmentally labeled Tm1 intermediate filaments also analyzed at 16°C (blue). Only a single contour is shown to clarify the peak positions in the two spectra. Residue type assignments are indicated based on random coil chemical shift values for CA and CB sites (*28*), or the average chemical shifts from the Biological Magnetic Resonance Data Bank for CG, CD, CE sites (*29*). We observed signals from every residue type in the tail domain outside of the region assigned to the immobilized core of the polymers, including the 6His tag and linker.

The two spectra have nearly identical peak positions for all sites. There are subtle differences for the positions of the signals in the histidine CA and CB, aspartic acid CA, asparagine CA, and leucine CA regions. Five extra signals were observed in the spectrum of the assembled intermediate filament sample and are indicated with asterisks in Fig. 3C. The only signals in the 6His-tag region not in the Tm1 tail domain are methionine, glutamine, tyrosine, and phenylalanine. It was not possible to unambiguously assign signals to these residues. Regardless, the spectra in Fig. 3C are consistent with both the His-tag and regions outside residues 384 to 411 as being highly dynamic in the polymer assemblies. For the intermediate filament assemblies, the high degree of similarity with regard to the spectrum of the Tm1 tail domain polymer assemblies indicates that there are nearly identical regions of structural disorder and dynamics in the two samples.

A TALOS-N prediction of backbone φ/ψ torsion angles based on the assigned ^13^C and ^15^N NMR chemical shifts from the CP-based spectra (*30*) is shown in Fig. 3D. The analysis uses a database of known protein structures to predict the torsion angles using amide ^15^N, ^13^CA, ^13^CB, and ^13^CO chemical shifts. Residues N387 to S392, N398 to A399, and G404 have β-strand secondary structure predictions and the extended regions of S386 to S392, S394 to E395, N398 to D401, G404 to N407, and A410 to S411 have backbone φ/ψ torsion angle predictions that are characteristic of extended β-strands. The zf-TEDOR measurements, the TALOS chemical shift analysis, and previous X-ray diffraction studies (*7*) confirm that Tm1 tail domain-only polymers adopt a cross-β structural conformation.

### Functional assays of deletion mutations in the LC tail domain of Tm1

In order to study the functional importance of the 69 residues specifying the Tm1 tail domain we prepared and analyzed a series of deletion mutants. Our mutations systematically deleted sequences starting at the C-terminus of the Tm1 tail domain, with individual mutants sequentially removing five additional residues. Ten deletion mutants were assayed to measure both the formation of cross-β polymers made from the isolated tail domain, as well as the formation of intermediate filaments made from the otherwise intact Tm1 polypeptide. Three changes in tail domain-only polymer formation were observed as a consequence of ever-greater deletion of C-terminal tail domain residues (Fig. 4). Crossing of a boundary between residues 425 and 430 yielded polymers that changed morphology from smooth to twisted. Crossing of a second boundary between residues 405 and 410 yielded polymers that changed morphology from twisted to spiky. Finally, crossing of a boundary between residues 390 and 395 yielded no cross-β polymers at all.

**Fig. 4.**
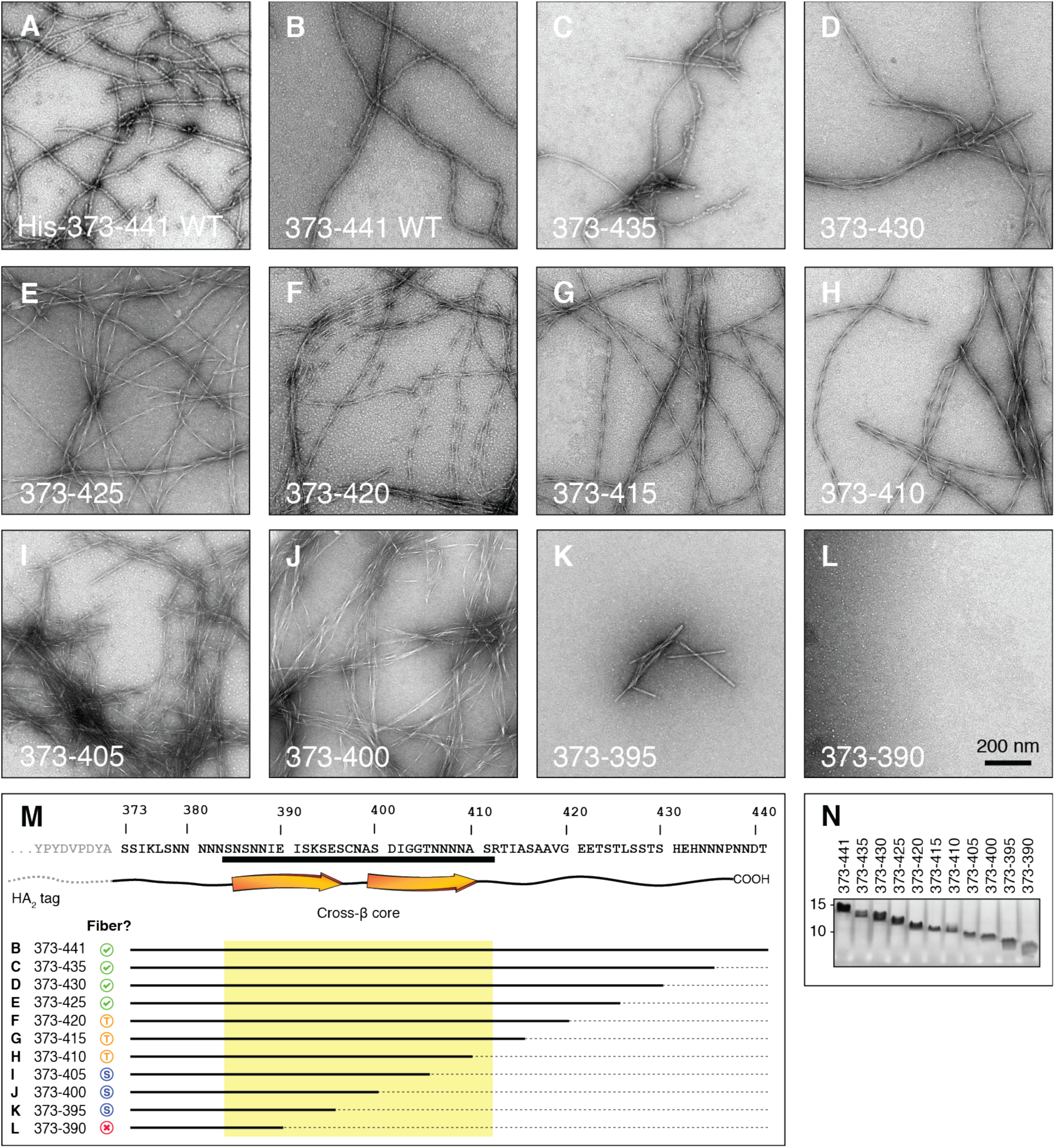
Functional assays of Tm1 tail domain-only polymer formation for ten C-terminal deletion mutants. Tm1 tail domain variants missing 6, 11, 16, 21, 26, 31, 36, 41, 46 or 51 residues from the C-terminus were expressed in bacterial cells, purified and incubated under conditions suitable for tail domain polymerization (see Materials and Methods). Panels A and B show intact tail domain polymers bearing an N-terminal His-tag (A) or an N-terminal HA_2_-tag (B). Variants missing up to 16 amino acids formed polymers similar to the intact HA_2_-tagged Tm1 tail domain (panels C-E). Variants missing between 21 and 31 amino acids formed distinctly twisted (T) polymers (panels F-H). Variants missing between 36 and 46 amino acids formed thin, spiky (S) polymers (panels I-K). The variant missing 51 C-terminal amino acids did not form polymers (panel L). All electron micrographs were taken at the same magnification (scale bar in panel L = 200 nm). Bottom left panel (M) shows Tm1 tail deletion mutants relative to the amino acid sequence of the tail domain and the location of the cross-β core determined in this work (orange arrows and yellow highlight). Bottom right panel (N) shows Coomassie brilliant-blue stained SDS polyacrylamide gel used to visualize purified Tm1 tail domain and ten C-terminal deletion mutants. Numbers to left of the SDS-PAGE gel designate migration positions of molecular weight markers at 15 and 10 kDa.

It was likewise observed that changes in Tm1 intermediate filament formation also suffered sequential impediments as a function of C-terminal deletion of the Tm1 tail domain (Fig. 5). Crossing of a boundary between residues 425 and 420 yielded proteins that changed filament morphology from normal to an aggregated, hairball configuration. Crossing of boundary between residues 405 and 400 yielded variant forms of the Tm1 protein that failed to assemble into intermediate filaments.

**Fig. 5.**
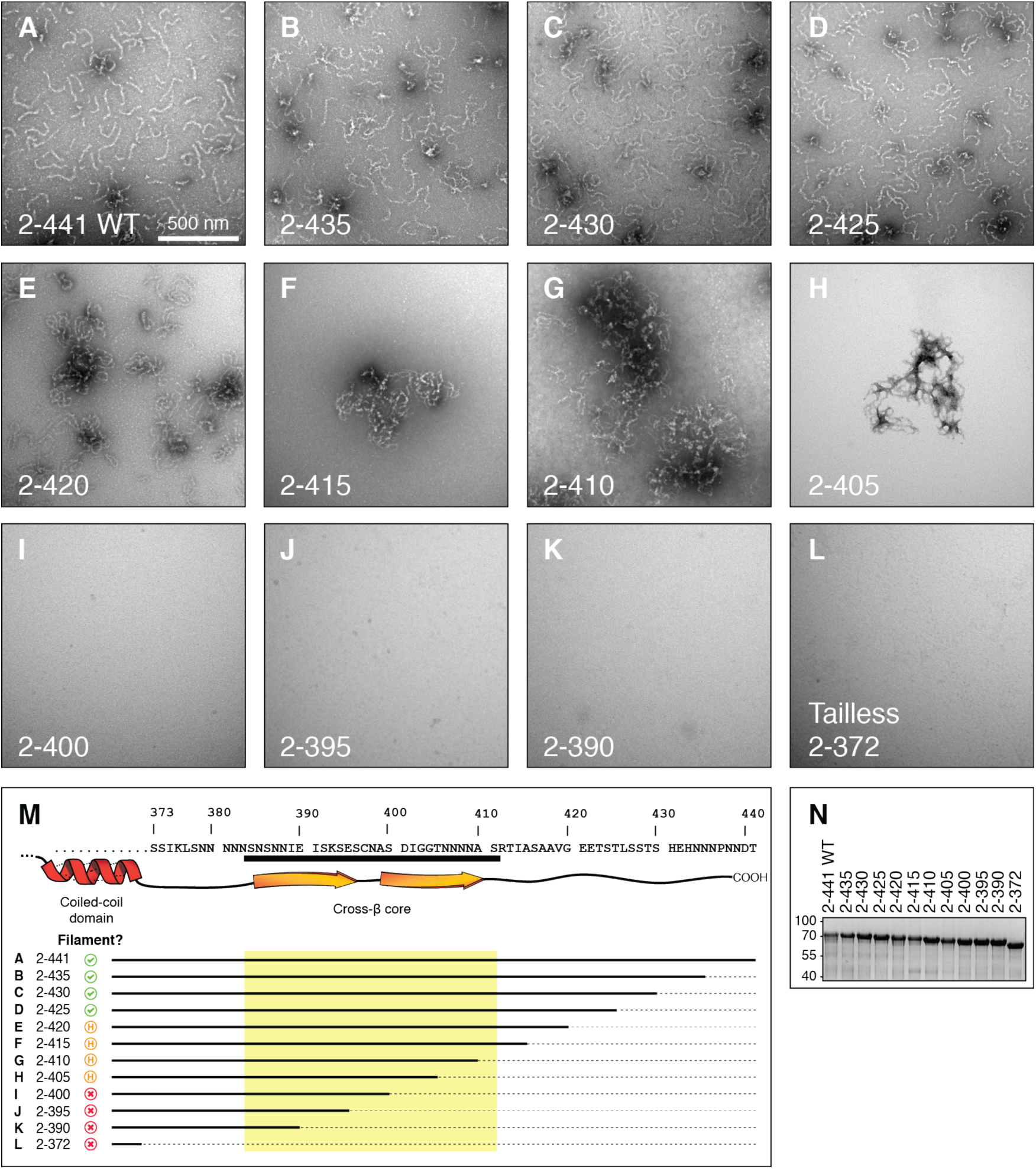
Functional assays of the ability of Tm1 C-terminal deletion mutants to assemble into intermediate filaments. Deletion variants of the otherwise full length Tm1 protein missing 6, 11, 16, 21, 26, 31, 36, 41, 46, 51 or 69 C-terminal residues were expressed in bacterial cells, purified and incubated under conditions allowing for the formation of intermediate filaments (see Materials and Methods). Tm1 variants missing up to 16 C-terminal residues of the protein formed intermediate filaments indistinguishable from those made from the intact protein (panels A-D). Variants missing between 21 and 36 C-terminal residues yielded aberrantly assembled filaments of a tangled or hairball-like morphology (panels E-H). Variants of the Tm1 protein missing 41 or more C-terminal residues failed to form intermediate filaments (panels I-L). Bottom left panel (M) shows Tm1 tail domain deletion mutants relative to the amino acid sequence of the tail domain, the location of the C-terminal end of the coiled-coil domain (red), and the location of the cross-β core determined in this work (orange arrows and yellow highlight). Bottom right panel (N) shows Coomassie brilliant-blue stained SDS polyacrylamide gel used to visualize purified samples of both intact (WT) Tm1 and eleven C-terminal deletion mutants. Numbers to the left of the SDS-PAGE gel designate migration position of molecular weight markers at 100, 70, 55 and 40 kDa.

## Discussion

Here we report studies of an intermediate filament produced in the oocytes of *Drosophila melanogaster*. Exons of the tropomyosin gene are alternatively spliced in oocytes so as to produce an unusual isoform of tropomyosin. Other than nuclear lamins, the Tm1 protein represents the only intermediate filament protein found in fruit flies. The central, coiled-coil domain of the Tm1 isoform of tropomyosin is flanked by low complexity sequences. Instead of interacting with troponin in a manner allowing for calcium-mediated regulation of muscle contraction, the C-terminal LC domain directs the Tm1 protein to assemble into intermediate filaments.

Why is proper deposition of germ granules at the posterior pole of *Drosophila* oocytes dependent upon Tm1-specified intermediate filaments? Visualization of a GFP-tagged form of Tm1 in living oocytes has shown that the protein becomes precisely restricted to the posterior tip of oocytes well before the completion of polar granule deposition (*8*). We speculate that assembled Tm1 intermediate filaments localized to the posterior pole of fly oocytes might constitute a Velcro-like landing pad for one or more of the constituent RNA binding proteins specifying polar granules.

In an architectural sense, assembled intermediate filaments contain repeating collars of low complexity domain sequences circumferentially displayed along their axial length (Supplementary Data, Fig. S1). In the case of vimentin intermediate filaments, we have observed that these LC domain collars represent repetitively organized binding sites for a GFP:FUS fusion protein, such that the latter protein can be iteratively bound to the filaments at 45 nm intervals as viewed by transmission electron microscopy (*15*). It is perhaps of importance that binding of the GFP:FUS protein to vimentin intermediate filaments is dependent upon the integrity of the LC tail domain of vimentin. As such, we speculate that Tm1 intermediate filaments, upon localization to the posterior tip of fly oocytes, may allow for the subsequent binding and organization of the germ granules themselves.

The experiments described in the present study have been focused on the C-terminal LC domain of the Tm1 protein, herein referred to as the tail domain. The Tm1 tail domain has already been shown to be required for Tm1 to assemble into intermediate filaments. Moreover, the isolated Tm1 tail domain has already been shown to form labile, cross-β polymers (*7*). The sole focus of the present study has been to determine whether the same structural forces leading to the formation of cross-β polymers observed in studies of the isolated Tm1 tail domain might also be involved in assembly of Tm1 intermediate filaments.

This central question has been approached via the use of ss-NMR spectroscopy. We have compared ss-NMR spectra observed from isotopically-labeled tail domain-only polymers with spectra derived from fully assembled intermediate filaments. In the latter case, we used intein chemistry to restrict isotopic labeling to only the tail domain of the intact Tm1 protein. The ^13^C/^15^N isotopic labels segmentally introduced into the intact Tm1 protein were identical to the labels introduced into tail domain-only polymers.

We highlight four observations from these studies. First, highly similar spectra diagnostic of structural order were observed in both tail domain-only cross-β polymers and segmentally labeled Tm1 intermediate filaments (Fig. 2). The portion of the Tm1 tail domain specifying these cross-β interactions was mapped to a region of 28 amino acids spanning residues 384 to 411 of the Tm1 polypeptide (Fig. 3). Second, evidence of robust molecular order within the tail domain was observed only upon cooling of the Tm1 intermediate filaments (Fig. 1). Third, roughly 30 amino acid residues were observed to exist in a state of distinct structural disorder in both the tail domain-only cross-β polymers and the fully assembled, segmentally-labeled intermediate filaments. Importantly, there was almost perfect overlap of the spectra reporting upon these disordered residues in the two structures (Fig. 3). Fourth, the observed φ/ψ torsion angles based upon ^13^C and 15^N^ chemical shifts for almost all residues within the structurally ordered tail domain of Tm1 were predictive of either β-strand secondary structure or extended β-strand-like conformation (Fig. 3).

Our parsimonious interpretation of these observations is that Tm1 intermediate filaments assemble via the formation of dynamically ordered cross-β interactions specified by the amino acid sequence of the C-terminal tail domain. When studied in the context of isolated, tail domain-only polymers, these cross-β interactions remain labile to disassembly, but are far more stable than those formed in the context of biologically relevant Tm1 intermediate filaments. This interpretation posits an unusual form of protein structure. We hereby offer that what is being observed in this study is a form of protein structure that retains an ordered ground state sufficiently precise to dictate both molecular order and functional specificity. Of curiosity, however, is the apparent fact that the amino acid residues specifying molecular order are in a state of continuous flux in and out of the structurally ordered state. We speculate that what we have observed in studies of the low complexity domain specifying function of the C-terminal tail domain of the Tm1 protein may be instructive of the thousands of LC domains operative in many other aspects of cell biology.

We close with emphasis on the fact that no more than one third of the amino acids of the Tm1 tail domain achieve any state of molecular order whether studied in either tail domain-only polymers or assembled intermediate filaments. As shown in Fig. 3C, precisely the same 30+ amino acid residues exist in a state of molecular disorder in both tail domain-only polymers and segmentally labeled intermediate filaments. That these disordered residues are likely to be of functional importance is supported by studies of systematically deleted variants of the Tm1 polypeptide. Deletion of only 20 residues from the C-terminus of the Tm1 protein, despite not impinging whatsoever upon the structurally ordered region of the tail domain, significantly impedes assembly of intermediate filaments and yields tail domain-only polymers having a distinctly twisted morphology (Figs. 4 and 5). That LC domains utilize no more than a modest fraction of their amino acid sequences to achieve transient structural order has also been reported for both the hnRNPA2 and fused in sarcoma (FUS) RNA binding proteins (*10, 24*). Why LC domains are reliant upon regions specifying both structural order and disorder remains an intriguing mystery.

## Acknowledgements

We thank Robert Tycko for thoughtful discussions regarding all aspects of the research described herein. SLM was supported by NIGMS grant 5R35GM13130358 as well as unrestricted funding provided by an anonymous donor. DTM was supported by faculty startup funding provided by the University of California at Davis. A portion of this work was performed at the National High Magnetic Field Laboratory, which is supported by the NSF Cooperative Agreement DMR-1644779 and the State of Florida. We also thank Dr. Ivan Hung for assistance with the NMR spectrometers at the Tallahasse, Florida facility. Experiments were planned by SLM, MK and DTM. VOS and LS cloned, mutagenized and expressed all recombinant proteins in both unlabeled and isotopically labeled forms. VOS assembled all tail domain-alone polymers and intermediate filaments formed from the Tm1 protein. RH and DTM recorded and interpreted all NMR spectra. SLM composed the manuscript with help from VS, MK and DTM. No authors of this manuscript have any form of competing interest. All data in this manuscript and supplementary materials are freely available.

## List of Supplementary Materials

Materials and Methods

Figures S1-S6

Tables S1-S5

References 31-33

