## Supplemental Materials for "Dynamic structural order of a low complexity domain facilitates assembly of intermediate filaments"

**Affiliations:**

**This PDF file includes:**

Materials and Methods
Figs. S1 to S6
Tables S1 to S5
References 31 to 33

### Materials and Methods:

**Protein expression and purification:** DNA fragments containing Tm1 coding sequence were produced by PCR and inserted into pHis-parallel vector as previously described (*7*). For production of DNA fragments encoding $Cfa_{GEP}$ intein sequence we used plasmids created by Stevens et al. (*19*). His-tagged recombinant proteins were expressed in *E. coli* BL21 DE3 strain. For production of unlabeled proteins, starter cultures were diluted 1:100 in LB medium supplemented with 100 mg/L ampicillin, grown at 37°C to $OD_{600}$ of 0.6-1.0, induced with 0.5 mM IPTG and incubated at 18°C overnight.

For production of universally-labeled Tm1 tail domain constructs, His-Tm1-373-441 and His-GFP-GDEVD-$Cfa^C$-Tm1-373-441 starter cultures were diluted 1:100 in M9 medium (12.8 g/L $Na_2HPO_4$, 3 g/L $KH_2PO_4$, 0.5 g/L NaCl, 1 g/L $NH_4Cl$ [$^{15}N$], 2 g/L glucose [$^{13}C$], 2 mM $MgSO_4$, 0.1 mM $CaCl_2$, 10 mg/L thiamine, 10 mg/L $FeSO_4$, 100 mg/L ampicillin), grown at 37°C to $OD_{600}$ of 0.6-1.0, induced with 0.5 mM IPTG and incubated at 18°C overnight. Cells were harvested by centrifugation at 5,000xG for 15 min, washed once with 20 mM Tris-HCl pH 7.5, 150 mM NaCl, and either stored at -80°C or used immediately for protein purification.

Bacterial pellets were resuspended in lysis buffer (50 mM Tris-HCl pH 7.5, 500 mM NaCl, 4 M urea, 1 mM TCEP, 1% Triton-X100, 1x EDTA-free protease inhibitor cocktail, Sigma) and sonicated. Lysates were cleared by ultracentrifugation at 100,000xG for 60 minutes. Cleared lysates were mixed with equilibrated Ni-NTA resin (Goldbio) and incubated for 15 minutes. The resin was then washed with wash buffer (20 mM Tris-HCl pH 7.5, 500 mM NaCl, 4 M urea, 1 mM TCEP, 20 mM imidazole, 0.1 mM PMSF), bound protein was eluted using wash buffer supplemented with 300 mM imidazole and concentrated in centrifugal filters with MWCO 3 kDa. In case of proteins not containing intein fragments, 1 mM TCEP was substituted with 20 mM β-mercaptoethanol. The C-terminal intein fragment His-GFP-GDEVD-$Cfa^C$-Tm1-373-441 was diluted 1:5 with caspase 3 buffer (20 mM Tris-HCl pH 7.5, 100 mM NaCl, 1 mM EDTA, 0.1% CHAPS, 10% glycerol, 10 mM DTT) and cleaved with 50 μg/ml caspase 3 at room temperature overnight. Cleaved GFP tag was removed by gel filtration in 20 mM Tris-HCl pH 7.5, 200 mM NaCl, 4 M guanidine hydrochloride, 1 mM TCEP on a Superdex 200 column.

**Ligation of intein fragments and purification of segmentally labeled full-length Tm1:** 15 mg (1.3 μmol) of $^{15}N/^{13}C$-labeled CfaC-Tm1-373-441 was mixed with 30 mg (0.5 μmol) unlabeled N-terminal fragment His-Tm1-2-372-CfaN in 3 ml denaturing buffer (20 mM Tris-HCl, 200 mM NaCl, 4 M guanidine hydrochloride). The reaction mixture was dialyzed against intein reaction buffer (40 mM K-phosphate buffer pH 7.2, 150 mM NaCl, 1 mM TCEP) overnight at room temperature. N-terminal fragment quantitatively converted to full-length segmentally labeled Tm1, and was separated from the excess of C-terminal fragment by fractionation on a MonoQ ion exchange column with 0-500 mM NaCl gradient in 20 mM Tris-HCl pH 7.5, 4 M urea, 5 mM β-mercaptoethanol. The product was additionally purified by gel filtration in 20 mM Tris-HCl pH 7.5, 200 mM NaCl, 4 M guanidine hydrochloride, 5 mM β-mercaptoethanol, concentrated and stored at -80°C.

**Purification of $HA_2$-tagged tail domain truncations:** His-GFP-GDEVD-$HA_2$-Tm1 tail domain truncation constructs were expressed, purified on Ni-NTA and cleaved with caspase 3 as described above. Cleaved HA-Tm1 tail domain fragments were purified on a HiTrap phenyl sepharose column with 1-0 M reverse ammonium sulfate gradient in 20 mM Tris-HCl pH 7.5, 4 M urea, 5 mM β-mercaptoethanol and concentrated in centrifugal filters with MWCO 3 kDa. Buffer was first exchanged to denaturing buffer (20 mM Tris-HCl pH 7.5, 200 mM NaCl, 6 M guanidine hydrochloride, 5 mM β-mercaptoethanol) and then to gelation buffer (20 mM Tris-HCl pH 7.5, 200 mM NaCl, 5 mM β-mercaptoethanol) on a HiPrep desalting column. Protein solutions were diluted to 20 μM with gelation buffer, stored overnight at room temperature and used for electron microscopy.

**Preparation of Tm1 tail domain polymers:** His-Tm1-373-441 protein was concentrated to 10 mg/ml (1 mM) and dialyzed against gelation buffer (20 mM Tris-HCl pH 7.5, 200 mM NaCl, 5 mM β-mercaptoethanol) overnight at room temperature. Dialyzed protein solution was centrifuged at 3000xG in a tabletop centrifuge to remove insoluble particles, sonicated, concentrated to 50 mg/ml and incubated at 4°C until uniform long polymers formed as monitored by electron microscopy.

**Assembly of Tm1 intermediate filaments:** Full-length Tm1 and truncation constructs dissolved in denaturing buffer (20 mM Tris-HCl, 200 mM NaCl, 4 M guanidine hydrochloride, 5 mM β-mercaptoethanol) were diluted to 20 μM and dialyzed stepwise against buffers containing decreasing concentrations of urea (5 mM Tris-HCl pH 8.6, 1 mM EDTA, 0.1 mM EGTA, 5 mM β-mercaptoethanol, 8 M – 4 M – 2 M – 0 M urea), with each step taking two hours. For complete removal of urea, protein solutions were additionally dialyzed against 0 M urea buffer overnight. Protein solutions were then mixed 1:1 with 2x filament assembly buffer (45 mM MES-Na pH 6.25, 100 mM NaCl), incubated at room temperature overnight and analyzed by electron microscopy.

**TEM imaging of Tm1 assemblies:** 10 μl of protein solutions was applied to carbon film grids (CF400-Cu, Electron Microscopy Sciences) for 1 min and removed by blotting. Grids were briefly washed with 5 mM EDTA pH 8.0, exposed to 2% uranyl acetate solution for 15 seconds and air-dried. Images were obtained with JEOL 1200EX or Tecnai Spirit electron microscopes.

**Solid State NMR Sample Packing:** Tm1 tail domain polymers were centrifuged at 150,000xG for 30 min and the supernatant discarded. The concentrated pellet was packed into a 3.2 mm thin-walled Varian style NMR rotor by centrifugation at 16,000xG for 2 h in a swinging bucket rotor with a homemade rotor holder, excess buffer removed with the tip of a laboratory wipe, and caps sealed with superglue gel. 1 mm teflon spacers were used to center the sample in the NMR rotor. Segmentally labeled Tm1 tail domain in intermediate filament assemblies were centrifuged at 150,000xG for 30 min and the supernatant discarded. Concentrated pellets from three centrifugations were combined and centrifuged at 4°C for 2 h at 365,000xG. The supernatant was again removed and the pellets packed into a 3.2 mm thin-walled Varian style NMR rotor by centrifugation at 17,000xG for 7.5 h, excess buffer removed with the tip of a laboratory wipe, and caps sealed with superglue gel. For the ether precipitated Tm1 tail domain sample, 0.5 ml of 3 mM His-tagged Tm1 tail domain in denaturing buffer (20 mM Tris-HCl pH 7.5, 6 M guanidine hydrochloride, 5 mM β-mercaptoethanol) was dialyzed against 2 L of ultrapure water for 16 h, exchanged for fresh ultrapure water and dialyzed for 8 h more. The precipitate was harvested, flash frozen in liquid nitrogen, and lyophilized for 16 h. The lyophilized protein was resuspended in 100 μl of trifluoroacetic acid and vortexed until the solution was clear. 1 ml of ice cold tertbutyl methyl ether was added to the solution, vortexed for 30 s, then centrifuged at 16,000xG for 20 min in a swinging bucket rotor precooled to 4°C. The supernatant was discarded and the pellet dried under a stream of nitrogen gas. The Tm1 tail domain powder was then packed into a 3.2 mm medium-walled Varian style NMR rotor with 1 mm spacers to center the sample in the rotor using a funnel and packing rod.

**Solid State NMR Measurements:** Solid state NMR measurements were performed on a 17.5 T superconducting magnet with a Varian InfinityPlus console and a BlackFox LLC triple resonance 3.2 mm MAS probe and on a 18.8 T superconducting magnet with a Bruker NEO console and a 3.2 mm MAS triple resonance probe built at the National High Magnetic Field Laboratory. All spectra are referenced using the upfield $^{13}C$ chemical shift of adamantane as 40.48 ppm (*31*). Sample temperatures were maintained using a stream of nitrogen gas and calibrated using a sample of potassium bromide (*32*). All experiments utilized 12 to 13 kHz MAS spinning. $^{1}H$ decoupling was either TPPM or SPINAL64 for CP based spectra or WALTZ-16 for INEPT based spectra. The CP ramp was 20% on $^{1}H$ during $^{1}H$-$^{13}C$ or $^{1}H$-$^{15}N$ CP and 5% on $^{15}N$ during $^{15}N$-$^{13}C$ SCP. See Table S5 for a complete list of NMR acquisition and processing parameters.

**Assignment Calculations and Torsion Angle Predictions:** Signal tables (Supplemental Tables S3 and S4) were generated from the 3D NCACX and 3D NCOCX spectra manually using the Sparky NMR visualization software (*33*). Uncertainties for each signal were assigned based on the signal to noise and resonance linewidth in each data set. Typical uncertainty values for $^{15}N$ sites were 0.4 and 0.3 for $^{13}C$ sites. Residue types were assigned based on the $^{13}C$ chemical shifts and the typical ranges for each amino acid from the Biological Magnetic Resonance Bank (*29*). The signal tables were checked for consistency with the 2D NCACX and 2D NCOCX spectra before using the simulated annealing monte carlo assignment algorithm MCASSIGN2B (*23*) to determine statistically significant sequence-specific assignments. All MCASSIGN2B calculations used the complete sequence of the 6His-tagged Tm1 tail domain-only construct and compared the observed amide $^{15}N$ and CA, CO, CB, and CG/CD $^{13}C$ chemical shifts.

In a first round of calculations, the ‘good’, ‘bad’, ‘edge’, and ‘used’ weights were ramped from 0 to 10, 10 to 60, 0 to 6, and 0 to 2 in 160 steps with 1 x $10^7$ iterations per step. An acceptance threshold was set to 0.001 so that the weights were ramped up in five steps to their final values if the number of accepted moves in the assignment algorithm became too small. 50 independent calculations were performed. In this first round, residues with assignments in 100% (*i.e.* 50 out of 50) of the calculations were S40 to S42, I45 to T61 (the NCACX signal for S52 was not assigned in 100% of the runs), and N65 to A66.

In a second round of calculations, these assignments were added to the input signal tables and the uncertainty of the $^{15}N$ chemical shift value was increased to 0.5 ppm for signal 23 in the NCACX signal table and for signal 16 in the NCOCX signal table. 50 individual calculations were run again using the same parameters as in the first round, but only 120 steps were used. Unique assignments were obtained in 100% of the second round calculations for the additional residues N43 to N44, S52 (the previously unassigned NCACX signal), N63 to N64, and S67.

For the third and final round of calculations, the signal tables were then reexamined and signals in the NCOCX table that represented self-correlations (*i.e.* $^{15}N_i$ to $^{13}C_i$ correlations within the same residue instead of $^{15}N_i$ to $^{13}C_{i-1}$ correlations) or signals that represented $^{13}C$-$^{13}C$ inter-residue magnetization transfers between neighboring amino acids were removed. Uncertainty values for the signal frequencies were increased for signal 11 in the NCACX table, and signals 18 and 27 in the NCOCX table. 40 steps were used in each of 50 additional calculations with all other parameters the same as in the previous two rounds of calculations. An additional assignment for residue N62 was made. This signal was not assigned to the same place in the sequence 100% of the time, but only one assignment had both an NCACX and NCOCX set of signals, which occurred in 18 of the 50 final calculations. Lists of the assigned and unassigned NCACX chemical shift values are presented in Supplemental Table S1 and S2, respectively.

For the torsion angle analysis, the assigned chemical shift values for the amide $^{15}N$ and $^{13}CA$, $^{13}CO$, and $^{13}CB$ chemical shifts were used as input to the TALOS-N torsion angle prediction program (*30*). The 'Strong' and 'Generous' predictions are plotted in Fig. 3D with error bars representing ± the estimated standard deviation of the prediction errors output by the TALOS-N program.

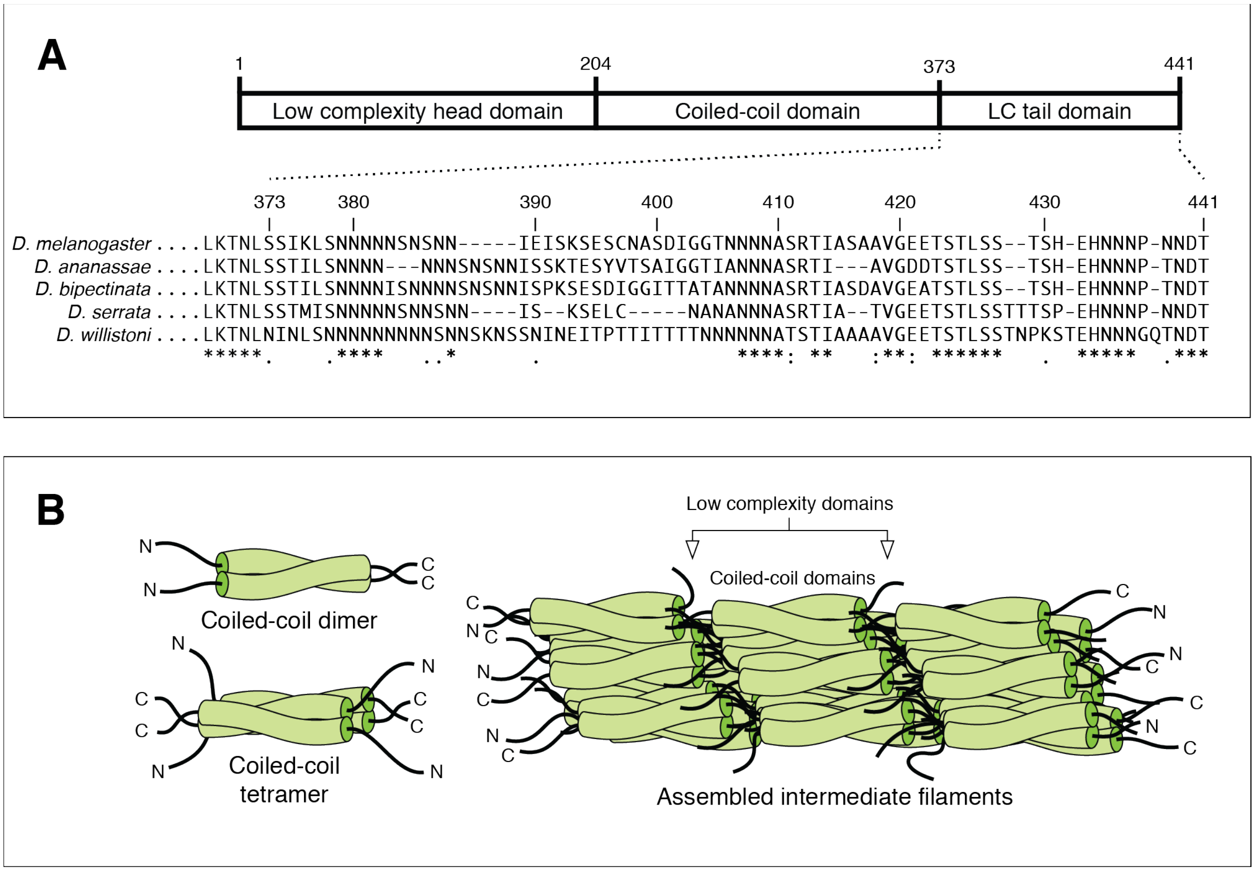


**Figure S1. Tm1 isoform of tropomyosin contains a low complexity tail domain required for assembly of intermediate filaments.** Top panel (A) shows domain organization of the Tm1 protein made in *Drosophila melanogaster* oocytes. Sequences of the C-terminal tail domains from individual species of the *Sophophora* sub-genus of *Drosophila* reveal three asparagine rich sub-domains within the five low complexity tail domains included in the alignment. Schematic diagram of Tm1 self-assembly is shown in the lower panel (B). Coiled-coil domains allow for formation of both parallel dimers and anti-parallel tetramers. Assembly of tetramers into intermediate filaments is dependent upon function of the low complexity tail domain **(***7***)**. Assembled intermediate filaments display a repetitive pattern of coiled-coil domains regularly interspersed by low complexity domains circumferentially positioned for self-interaction.

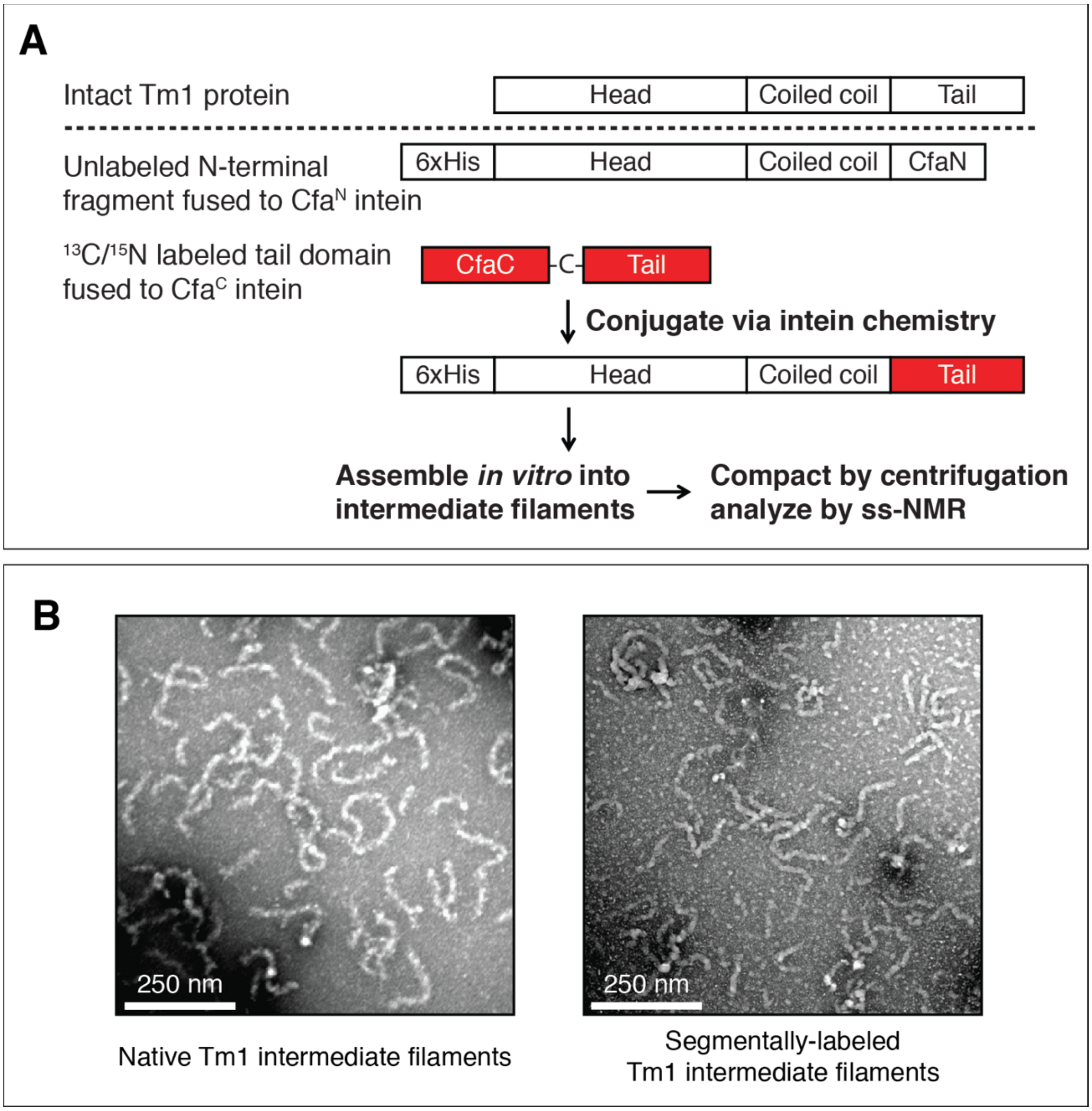


**Figure S2. Segmental isotope labeling of the Tm1 tail domain and its assembly into mature intermediate filaments.** Top panel (A) shows methods used to express a 6xHis-tagged form of Tm1 containing its low complexity head and coiled-coil domains linked to the Cfa$^N$ intein segment. The Tm1 tail domain linked to the Cfa$^C$ intein segment was labeled with $^{13}C/^{15}N$ isotopes (Materials and Methods). The two proteins were chemically conjugated, thus allowing removal of the Cfa$^N$ and Cfa$^C$ intein segments and fusion of the Tm1 tail domain to the otherwise intact Tm1 protein. Engineering of segmentally labeled Tm1 yielded a protein bearing a single difference, asparagine 371 changed to cysteine, relative to the native Tm1 protein. The intein-ligated, segmentally labeled protein was purified and incubated under conditions allowing formation of intermediate filaments. As shown in panel B, native and segmentally labeled Tm1 proteins assembled into intermediate filaments indistinguishable in transmission electron microscope images of negatively stained samples. Scale bars in both micrographs = 250 nm.

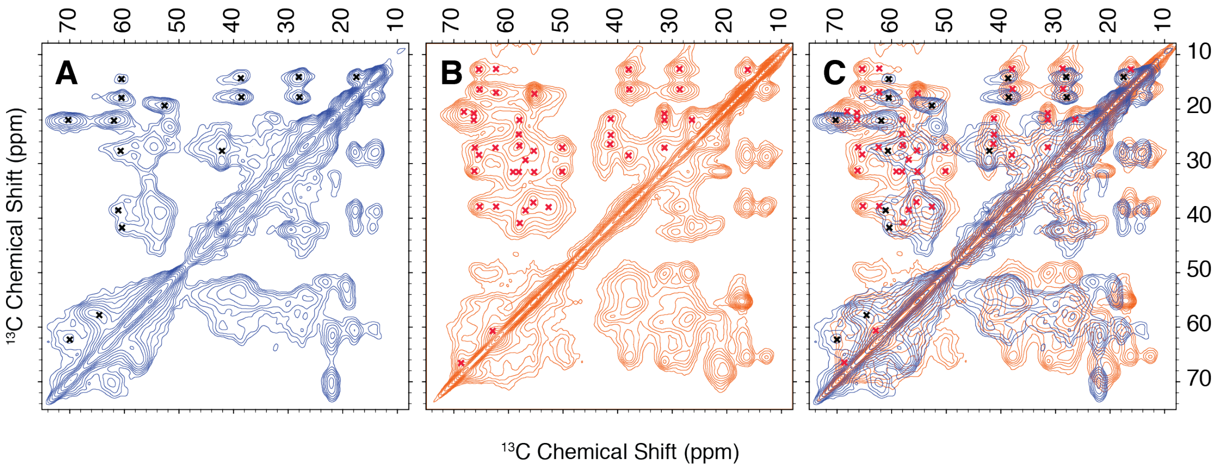


**Figure S3. Cross polarization-based solid state NMR spectra of the Tm1 tail domain.** (A) The $^{13}$C-$^{13}$C CP-DARR spectrum recorded at −19°C of the $^{13}$C/$^{15}$N segmentally labeled Tm1 tail domain that was assembled into mature intermediate filaments is shown in blue contours. (B) The $^{13}$C-$^{13}$C CP-DARR spectrum of the $^{13}$C/$^{15}$N labeled ether precipitated Tm1 tail domain recorded at 16°C is shown in orange contours. The X-marks displayed on the spectra indicate the center positions of the signals in the data recorded from the segmentally labeled full-length Tm1 filaments (black) and ether precipitated Tm1 tail domain (red). Panel C shows overlay of the two spectra. No significant overlap was observed between these two spectra. The contours are drawn with a factor of 1.4 between successive levels.

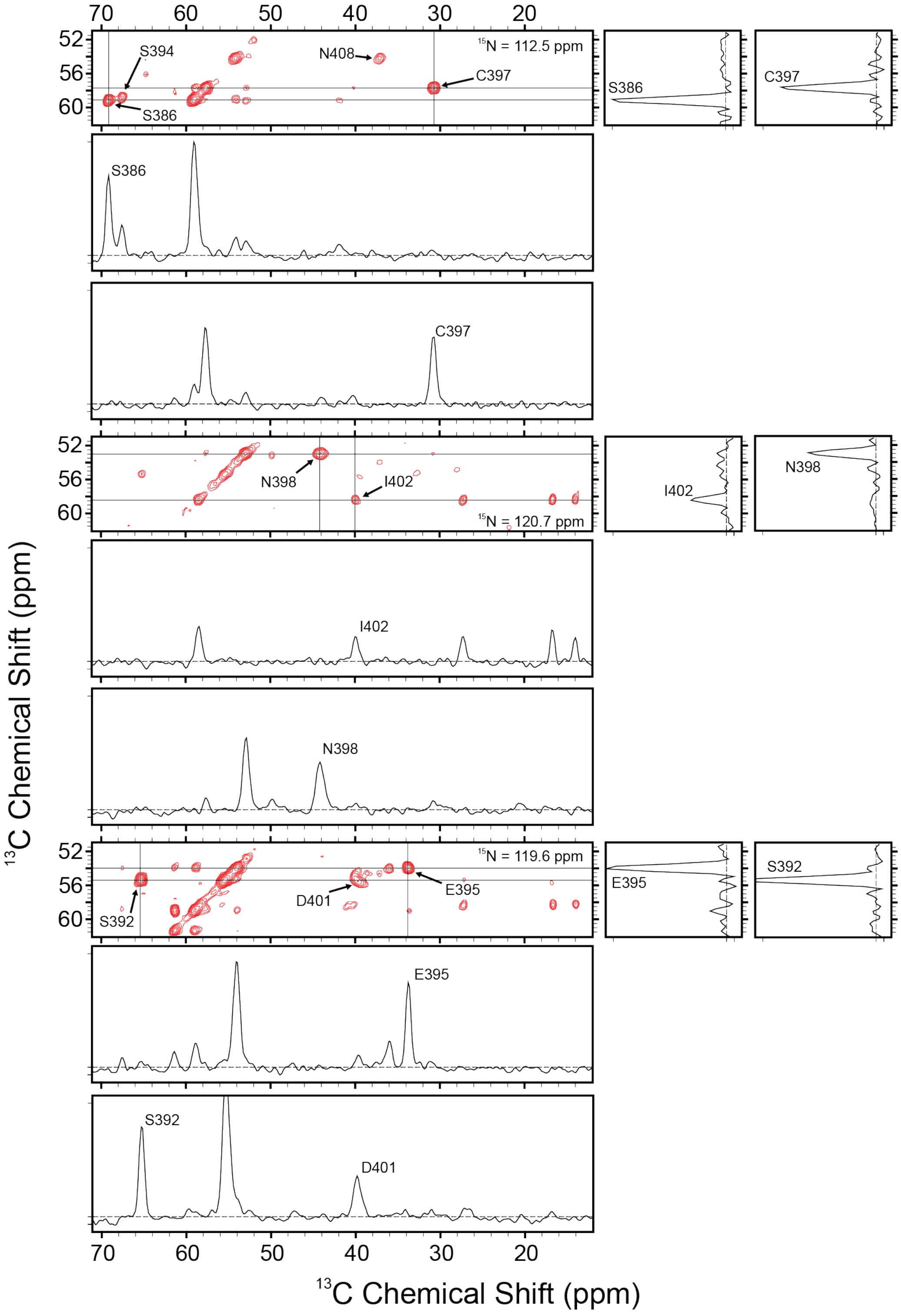


**Figure S4. Signal-to-noise in the 3D NCACX solid state NMR spectra of Tm1 tail domain-only polymers.** Three 2D slices from the cross polarization-based 3D NCACX spectrum of the $^{13}C/^{15}N$-labeled Tm1 tail domain-only polymer sample recorded at 16°C are shown with red contours drawn with a factor of 1.4 between successive levels. For each 2D slice, the two horizontal and vertical lines indicate the location of extracted 1D slices displayed below and to the right of the main panel. The residue numbers indicated in the plot are derived from sequence specific assignments determined in this work and refer to the numbering in the full-length Tm1 protein.

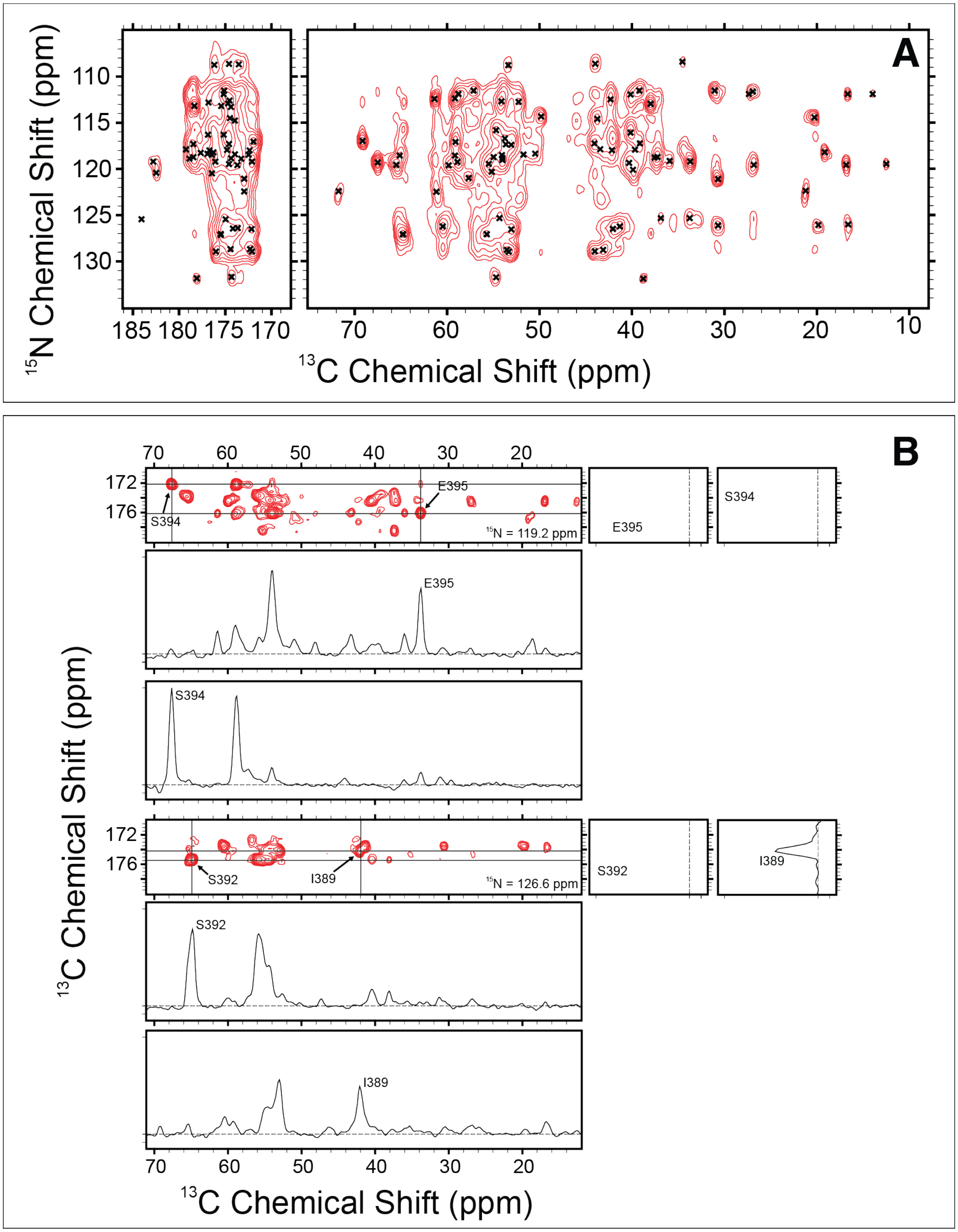


**Figure S5. 2D and 3D NCOCX solid state NMR spectra of the Tm1 tail domain-only polymers.** Top panel (A) shows a 2D cross polarization-based NCOCX spectrum of the $^{13}$C/$^{15}$N-labeled Tm1 tail domain polymers recorded at 16°C and drawn with a factor of 1.4 between contour levels. Black X's indicate the positions of signals determined from the 3D version of this spectrum. Bottom panel (B) shows two 2D slices from the cross polarization-based 3D NCOCX spectrum of the $^{13}$C/$^{15}$N-labeled Tm1 tail domain polymer sample recorded at

16°C and drawn with a factor of 1.4 between successive levels.  For each 2D slice, the two horizontal and vertical lines indicate the location of the 1D cross sections displayed below and to the right of the main panel.

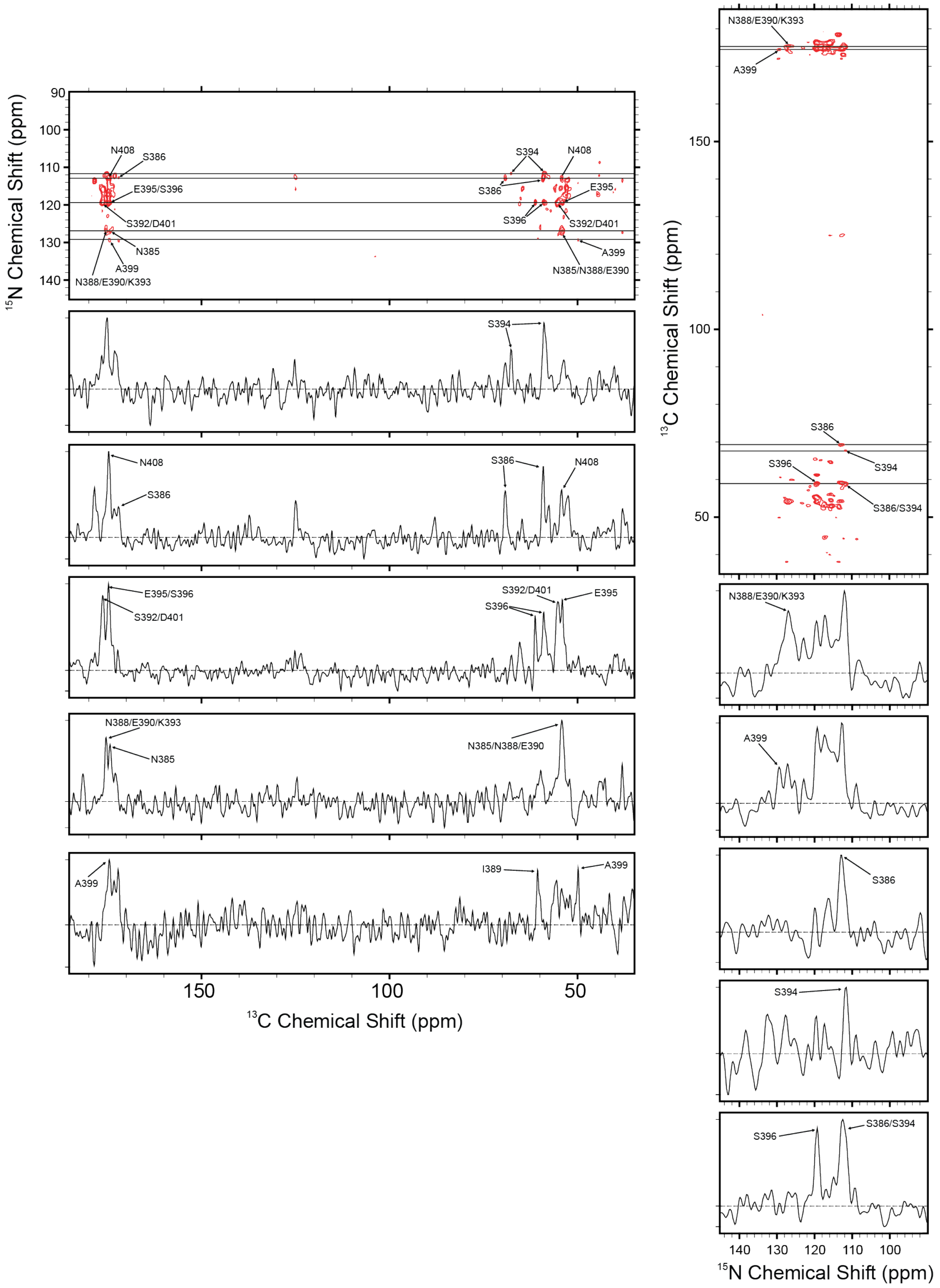


**Figure S6. Signal-to-noise in the 2D zf-TEDOR NMR spectra of Tm1 tail domain polymers.** Top left column is the 2D zf-TEDOR spectrum of the Tm1 tail domain-only polymers assembled from molecules of which 50% were labeled with $^{13}C/^{14}N$ and 50% were labeled with $^{12}C/^{15}N$. Horizontal lines drawn on this spectrum indicate the locations from which the series of 1D slices displayed below the spectrum were extracted. 1D slices are arranged in the same vertical order as the horizontal lines. The residue numbers indicated in the spectrum correspond to the numbering in the full length Tm1 protein and are derived from sequence specific assignments determined in this work. Right column contains the same 2D zf-TEDOR spectrum as the left column but rotated clockwise by 90°. As in the left column, the horizontal lines indicate the locations where 1D slices were extracted and displayed in the same vertical order as the horizontal lines. Residue numbers in this column also correspond to the residue numbering in the full length Tm1 protein and are derived from sequence specific assignments.

**Table S1.** Assigned NMR chemical shifts from the NCACX data.

| Assigned Signals | | | | | | | | | |
|---|---|---|---|---|---|---|---|---|---|
| **Residue Number** | **Residue Type** | **NH** | **CO** | **CA** | **CB** | **CG1** | **CG2** | **CD** | **S/N[1]** |
| 384 | S | 115.5 | 175.2 | 56.0 | 64.6 | | | | 56 |
| 385 | N | 127.2 | 174.3 | 54.2 | 38.0 | 178.3 | | | 43 |
| 386 | S | 112.4 | 172.0 | 59.1 | 69.2 | | | | 49 |
| 387 | N | 116.9 | 174.4 | 52.9 | 42.0 | 172 | | | 38 |
| 388 | N | 126.7 | 172.2 | 53.7 | 43.0 | 175.1 | | | 24 |
| 389 | I | 128.6 | 173.4 | 60.5 | 41.2 | 30.6 | 19.7 | 16.5 | 25 |
| 390 | E | 126.8 | 174.7 | 54.2 | 33.7 | 37.1 | | 184.0 | 18 |
| 391 | I | 125.4 | 174.4 | 59.8 | 40.5 | 26.8 | 16.8 | 12.5 | 21 |
| 392 | S | 119.6 | 175.2 | 55.3 | 65.2 | | | | 55 |
| 393 | K | 127.1 | 174.9 | 57.0 | 31.1 | 26.6 | | 39.2 | 14 |
| 394 | S | 111.5 | 172.1 | 58.8 | 67.5 | | | | 58 |
| 395 | E | 119.3 | 176.0 | 54.0 | 33.7 | 36.0 | 183.1 | | 53 |
| 396 | S | 119.1 | 174.5 | 59.0 | 61.3 | | | | 56 |
| 397 | C | 112.5 | 172.8 | 57.7 | 30.7 | | | | 41 |
| 398 | N | 120.9 | 172.0 | 53.0 | 44.1 | 176 | | | 29 |
| 399 | A | 129.0 | 174.4 | 49.8 | 20.3 | | | | 45 |
| 400 | S | 114.4 | 173.7 | 55.7 | 65.6 | | | | 33 |
| 401 | D | 119.5 | 175.9 | 55.4 | 39.7 | 182.5 | | | NR |
| 402 | I | 120.3 | 175.0 | 58.4 | 39.9 | 27.2 | 16.6 | 14.0 | 15 |
| 403 | G | 111.9 | 173.2 | 44.0 | | | | | -- |
| 404 | G | 108.6 | 173.7 | 43.8 | | | | | -- |
| 405 | T | 114.7 | 173.0 | 61.0 | 71.8 | | 21.4 | | 22 |
| 406 | N | 122.8 | 174.6 | 54.0 | 39.0 | 178.7 | | | 14 |
| 407 | N | 117.6 | 174.7 | 52.3 | 42.3 | 176.7 | | | 27 |
| 408 | N | 112.6 | 174.4 | 54.1 | 37.1 | 178.7 | | | 16 |
| 409 | N | 118.5 | 174.6 | 54.8 | 38.8 | 178.2 | | | 17 |
| 410 | A | 131.9 | 177.0 | 50.3 | 19.2 | | | | 28 |
| 411 | S | 118.3 | 174.3 | 59.1 | 65.1 | | | | 20 |

[1]Signal to noise (S/N) is the value reported by the Sparky NMR viewing program (*33*) for the CB resonance. NR indicates the residue was not resolved. Glycine residues have no CB atom.

**Table S2.** Unassigned NMR chemical shifts from the NCACX data.

| **Unassigned Signals** | | | | | | | | |
|---|---|---|---|---|---|---|---|---|
| **Residue Type** | **NH** | **CO** | **CA** | **CB** | **CG1** | **CG2** | **CD** | **S/N** |
| A | 126.1 | 175.6 | 50.9 | 18.6 | | | | 9 |
| T | 125.5 | 172.7 | 62.0 | 69.0 | | 22.0 | | 6 |
| A | 123.6 | 175.1 | 50.1 | 20.1 | | | | 13 |
| HMKRQECND | 123.4 | 176.0 | 53.4 | 34.4 | | | | 12 |
| N | 122.7 | 173.3 | 52.0 | 40.2 | 178.2 | | | 9 |
| T | 122.3 | 172.5 | 61.3 | 71.4 | | 21.1 | | 9 |
| N | 119.0 | 174.6 | 54.9 | 40.1 | 175.9 | | | NR |
| N | 117.4 | 173.2 | 55.2 | 37.4 | 178.3 | | | 17 |
| N | 117.1 | 173.4 | 53.5 | 39.2 | 178.8 | | | 20 |
| N | 116.3 | 174.5 | 52.9 | 34.3 | 176.1 | | | 11 |
| N | 115.5 | 175.8 | 51.2 | 42.2 | 177.1 | | | 11 |

[1]Signal to noise (S/N) is the value reported by the Sparky NMR viewing program (*33*) for the CB resonance. NR indicates the residue was not resolved.

**Table S3.** NCACX signal table used in the MCASSIGN calculations.[1]

| NH | CO | CA | CB | CX | NH Uncer. | CO Uncer. | CA Uncer. | CB Uncer. | CX Uncer. | Possible Amino Acid Types |
|---|---|---|---|---|---|---|---|---|---|---|
| 131.9 | 177.0 | 50.3 | 19.2 | 1111.1 | 0.4 | 0.3 | 0.3 | 0.3 | 0.3 | A |
| 129.0 | 174.4 | 49.8 | 20.3 | 1111.1 | 0.4 | 0.3 | 0.3 | 0.3 | 0.3 | A |
| 128.6 | 173.4 | 60.5 | 41.2 | 30.6 | 0.4 | 0.3 | 0.3 | 0.3 | 0.3 | I |
| 127.2 | 174.3 | 54.2 | 38.0 | 178.3 | 0.4 | 0.7 | 0.3 | 0.3 | 0.3 | ND |
| 127.1 | 174.9 | 57.0 | 31.1 | 26.6 | 0.4 | 0.3 | 0.3 | 0.3 | 0.3 | KR |
| 126.7 | 172.2 | 53.7 | 43.0 | 175.1 | 0.4 | 0.3 | 0.3 | 0.3 | 0.7 | ND |
| 126.8 | 174.7 | 54.2 | 33.7 | 37.1 | 0.4 | 0.5 | 0.3 | 0.4 | 0.4 | EQ |
| 125.4 | 174.4 | 59.8 | 40.5 | 26.8 | 0.4 | 0.3 | 0.3 | 0.3 | 0.3 | I |
| 126.1 | 175.6 | 50.9 | 18.6 | 1111.1 | 0.4 | 0.4 | 0.3 | 0.3 | 0.3 | A |
| 125.5 | 172.7 | 62.0 | 69.0 | 22.0 | 0.4 | 0.3 | 0.3 | 0.3 | 0.3 | T |
| 123.6 | 175.1 | 50.1 | 20.1 | 1111.1 | 0.4 | 0.4 | 0.3 | 0.3 | 0.3 | A |
| 123.4 | 176.0 | 53.4 | 34.4 | 1111.1 | 0.4 | 0.3 | 0.3 | 0.3 | 0.3 | HMKRQECND |
| 122.8 | 174.6 | 54.0 | 39.0 | 178.7 | 0.4 | 0.3 | 0.3 | 0.3 | 0.3 | ND |
| 122.7 | 173.3 | 52.0 | 40.2 | 178.2 | 0.4 | 0.3 | 0.3 | 0.3 | 0.3 | ND |
| 122.3 | 172.5 | 61.3 | 71.4 | 21.1 | 0.4 | 0.3 | 0.3 | 0.3 | 0.3 | T |
| 122.1 | 173.9 | 53.8 | 37.1 | 1111.1 | 0.4 | 0.3 | 0.3 | 0.3 | 0.3 | ND |
| 120.9 | 172.0 | 53.0 | 44.1 | 176 | 0.4 | 0.3 | 0.3 | 0.3 | 0.3 | ND |
| 120.3 | 175.0 | 58.4 | 39.9 | 27.2 | 0.4 | 0.3 | 0.3 | 0.3 | 0.3 | I |
| 119.6 | 175.2 | 55.3 | 65.2 | 1111.1 | 0.4 | 0.7 | 0.3 | 0.3 | 0.3 | S |
| 119.5 | 175.9 | 55.4 | 39.7 | 182.5 | 0.4 | 0.7 | 0.3 | 0.3 | 0.3 | ND |
| 119.3 | 176.0 | 54.0 | 33.7 | 36.0 | 0.4 | 0.3 | 0.3 | 0.3 | 0.3 | EQ |
| 119.1 | 174.5 | 59.0 | 61.3 | 1111.1 | 0.4 | 0.3 | 0.3 | 0.3 | 0.3 | S |
| 118.3 | 174.3 | 59.1 | 65.1 | 1111.1 | 0.5 | 0.3 | 0.3 | 0.3 | 0.3 | S |
| 119.0 | 174.6 | 54.9 | 40.1 | 175.9 | 0.4 | 0.6 | 0.3 | 0.3 | 0.6 | ND |
| 119.0 | 175.9 | 54.9 | 40.1 | 174.6 | 0.4 | 0.6 | 0.3 | 0.3 | 0.6 | ND |
| 118.5 | 174.6 | 54.8 | 38.8 | 178.2 | 0.4 | 0.5 | 0.3 | 0.3 | 0.3 | ND |
| 117.4 | 173.2 | 55.2 | 37.4 | 178.3 | 0.4 | 0.3 | 0.3 | 0.3 | 0.3 | ND |
| 117.1 | 173.4 | 53.5 | 39.2 | 178.8 | 0.4 | 0.4 | 0.3 | 0.3 | 0.3 | ND |
| 117.6 | 174.7 | 52.3 | 42.3 | 176.7 | 0.4 | 0.4 | 0.3 | 0.3 | 0.3 | ND |
| 116.9 | 174.4 | 52.9 | 42.0 | 172.0 | 0.4 | 0.5 | 0.4 | 0.3 | 0.3 | ND |
| 116.3 | 174.5 | 52.9 | 34.3 | 176.1 | 0.4 | 0.4 | 0.3 | 0.3 | 0.3 | ND |
| 115.5 | 175.2 | 56.0 | 64.6 | 1111.1 | 0.4 | 0.3 | 0.3 | 0.3 | 0.3 | S |
| 115.5 | 175.8 | 51.2 | 42.2 | 177.1 | 0.4 | 0.3 | 0.3 | 0.3 | 0.3 | ND |
| 114.7 | 173.0 | 61.0 | 71.8 | 21.4 | 0.4 | 0.3 | 0.3 | 0.3 | 0.3 | T |
| 114.4 | 173.7 | 55.7 | 65.6 | 1111.1 | 0.4 | 0.3 | 0.3 | 0.3 | 0.3 | S |
| 112.4 | 172.0 | 59.1 | 69.2 | 1111.1 | 0.4 | 0.3 | 0.3 | 0.3 | 0.3 | S |
| 112.5 | 172.8 | 57.7 | 30.7 | 1111.1 | 0.4 | 0.3 | 0.3 | 0.3 | 0.3 | HMKRQECND |
| 112.6 | 174.4 | 54.1 | 37.1 | 178.7 | 0.4 | 0.3 | 0.3 | 0.3 | 0.3 | ND |
| 111.5 | 172.1 | 58.8 | 67.5 | 1111.1 | 0.4 | 0.3 | 0.3 | 0.3 | 0.3 | S |
| 111.9 | 173.2 | 44.0 | 1111.1 | 1111.1 | 0.4 | 0.3 | 0.3 | 0.3 | 0.3 | G |
| 108.6 | 173.7 | 43.8 | 1111.1 | 1111.1 | 0.4 | 0.3 | 0.3 | 0.3 | 0.3 | G |

[1]A value of '1111.1' is used by the MCASSIGN program to indicate there is no observed signal for a particular atom.

**Table S4.** NCOCX signal table used in the MCASSIGN calculations.[1]

| NH | CO | CA | CB | CX | NH Uncer. | CO Uncer. | CA Uncer. | CB Uncer. | CX Uncer. | Possible Amino Acid Types |
|---|---|---|---|---|---|---|---|---|---|---|
| 132.1 | 174.4 | 54.8 | 38.8 | 178.1 | 0.4 | 0.3 | 0.3 | 0.3 | 0.3 | ND |
| 129.1 | 172.1 | 53.3 | 44.1 | 176.1 | 0.4 | 0.3 | 0.3 | 0.3 | 0.3 | ND |
| 128.8 | 172.3 | 53.4 | 43.1 | 174.4 | 0.4 | 0.3 | 0.3 | 0.3 | 0.3 | ND |
| 127.1 | 175.4 | 55.7 | 64.8 | 1111.1 | 0.4 | 0.3 | 0.5 | 0.3 | 0.3 | S |
| 127.1 | 175.4 | 55.7 | 64.8 | 1111.1 | 0.4 | 0.3 | 0.5 | 0.3 | 0.3 | S |
| 126.6 | 174.1 | 53.1 | 42.0 | 172.1 | 0.4 | 0.3 | 0.3 | 0.3 | 0.3 | ND |
| 126.4 | 173.5 | 60.6 | 41.3 | 30.6 | 0.4 | 0.3 | 0.3 | 0.3 | 0.3 | I |
| 125.4 | 175.1 | 54.4 | 33.8 | 37.0 | 0.4 | 0.3 | 0.4 | 0.3 | 0.3 | EQ |
| 122.4 | 172.9 | 61.2 | 71.8 | 21.2 | 0.4 | 0.3 | 0.3 | 0.3 | 0.3 | T |
| 121.1 | 172.9 | 57.7 | 30.8 | 1111.1 | 0.4 | 0.3 | 0.3 | 0.3 | 0.3 | CEQHKMR |
| 120.5 | 176.5 | 55.5 | 39.8 | 182.5 | 0.4 | 0.3 | 0.3 | 0.3 | 0.3 | ND |
| 119.5 | 172.1 | 58.8 | 67.6 | 1111.1 | 0.4 | 0.3 | 0.3 | 0.3 | 0.3 | S |
| 119.5 | 173.7 | 55.7 | 65.6 | 1111.1 | 0.4 | 0.3 | 0.3 | 0.3 | 0.3 | S |
| 119.7 | 174.5 | 59.9 | 40.5 | 26.9 | 0.4 | 0.3 | 0.3 | 0.3 | 0.3 | I |
| 119.1 | 176.1 | 54.0 | 33.8 | 36.0 | 0.4 | 0.3 | 0.3 | 0.3 | 0.3 | EQ |
| 118.9 | 176.8 | 50.5 | 19.2 | 1111.1 | 0.5 | 0.3 | 0.3 | 0.3 | 0.3 | A |
| 118.8 | 174.4 | 54.1 | 37.1 | 178.8 | 0.4 | 0.3 | 0.3 | 0.3 | 0.3 | ND |
| 118.7 | 173.3 | 55.3 | 37.4 | 178.4 | 0.4 | 0.3 | 0.3 | 0.3 | 0.3 | ND |
| 118.2 | 174.3 | 54.9 | 34.5 | 176.1 | 0.4 | 0.3 | 2 | 0.3 | 0.3 | ND |
| 118.5 | 173.9 | 59.2 | 65.1 | 1111.1 | 0.4 | 0.3 | 0.3 | 0.3 | 0.3 | S |
| 117.2 | 174.7 | 54.0 | 39.0 | 178.5 | 0.5 | 0.3 | 1.2 | 0.4 | 0.4 | ND |
| 117.0 | 171.9 | 59.2 | 69.3 | 1111.1 | 0.4 | 0.3 | 0.3 | 0.3 | 0.3 | S |
| 116.9 | 172.9 | 57.3 | 31.1 | 1111.1 | 0.4 | 0.3 | 0.3 | 0.3 | 0.3 | CEQHKMR |
| 116.3 | 175.7 | 51.3 | 42.4 | 177.3 | 0.4 | 0.3 | 0.3 | 0.3 | 0.3 | ND |
| 115.9 | 174.8 | 55.0 | 40.2 | 176.4 | 0.4 | 0.3 | 0.3 | 0.3 | 0.3 | ND |
| 114.8 | 173.9 | 43.9 | 1111.1 | 1111.1 | 0.4 | 0.3 | 0.3 | 0.3 | 0.3 | G |
| 114.5 | 174.6 | 49.9 | 20.4 | 1111.1 | 0.4 | 0.4 | 0.3 | 0.3 | 0.3 | A |
| 112.9 | 174.9 | 52.4 | 42.3 | 176.7 | 0.4 | 0.3 | 0.3 | 0.3 | 0.4 | ND |
| 112.6 | 174.4 | 54.3 | 38.1 | 178.4 | 0.4 | 0.3 | 0.3 | 0.3 | 0.3 | ND |
| 112.6 | 174.6 | 59.0 | 61.4 | 1111.1 | 0.4 | 0.3 | 0.3 | 0.3 | 0.3 | S |
| 111.8 | 175.1 | 58.4 | 40.1 | 27.2 | 0.4 | 0.3 | 0.3 | 0.3 | 0.3 | I |
| 111.8 | 173.6 | 53.6 | 39.3 | 178.9 | 0.4 | 0.4 | 0.3 | 0.3 | 0.4 | ND |
| 111.6 | 175.1 | 57.2 | 31.0 | 26.6 | 0.4 | 0.3 | 0.3 | 0.3 | 0.3 | KR |
| 108.4 | 176.2 | 53.5 | 34.5 | 1111.1 | 0.4 | 0.3 | 0.3 | 0.3 | 0.3 | HMKRQECND |
| 108.6 | 173.5 | 44.1 | 1111.1 | 1111.1 | 0.4 | 0.3 | 0.3 | 0.3 | 0.3 | G |

[1]A value of '1111.1' is used by the MCASSIGN program to indicate there is no observed signal for a particular atom.

**Table S5.** NMR experiments and parameter values for the data presented in this paper.

| Sample | Spectrum | NMR Parameters* | Exp. Time | Processing Parameters** |
|---|---|---|---|---|
| Uniform $^{13}C/^{15}N$ Tm1 Tail Domain-Only Polymers | 1D $^{13}C$ CP | $B_0$=18.8 T; $\nu_{MAS}$=13 kHz; $\tau_{\pi/2 1H}$=2.5 μs; $\nu_{CP1H}$=78 kHz; $\nu_{CP13C}$=63 kHz; $\tau_{CP}$ =1.0 ms; $\tau_{\pi SPINAL}$=4.9 μs; $\nu_{dec}$=100 kHz; $\tau_{acq}$=5.12 ms; $\tau_{dwell}$=5 μs; $n_{scan}$=1024; $\tau_{recycle}$=2 s; T=16 ºC; | 0.6 h | $t_1$: 90 Hz |
| Uniform $^{13}C/^{15}N$ Tm1 Tail Domain-Only Polymers | 2D $^{13}C$-$^{13}C$ CP-DARR | $B_0$=17.5 T; $\nu_{MAS}$=12 kHz; $\tau_{\pi/2 1H}$=4 μs; $\tau_{\pi/2 13C}$=5 μs; $\nu_{CP1H}$=52 kHz; $\nu_{CP13C}$=38 kHz; $\tau_{CP}$ =1.5 ms; $\tau_{\pi TPPM}$=5.6 μs; $\nu_{dec}$=83 kHz; $\tau_{DARR}$= 50 ms; $\nu_{DARR}$=12 kHz; $\Delta t_1$=22.4 μs; $\tau_{t1}$=4.48 ms; $t_{1MODE}$=States; $\tau_{acq}$=7.68 ms; $\tau_{dwell}$=15 μs; $n_{scan}$=256; $\tau_{recycle}$=1.5 s; T=16 ºC; | 42.7 h | $t_1$: 90 Hz/312 pts<br>$t_2$: 90 Hz |
| Uniform $^{13}C/^{15}N$ Tm1 Tail Domain-Only Polymers | 1D $^{13}C$ INEPT | $B_0$=18.8 T; $\nu_{MAS}$=13 kHz; $\tau_{\pi/2 1H}$=10 μs; $\tau_{\pi/2 13C}$=8.3 μs; $\tau_J$=1.06 ms; $\nu_{dec}$=25 kHz; $\tau_{acq}$=10.2 ms; $\tau_{dwell}$=10 μs; $n_{scan}$=1024; $\tau_{recycle}$=1 s; T=16 ºC; | 0.3 h | $t_1$: 50 Hz |
| Uniform $^{13}C/^{15}N$ Tm1 Tail Domain-Only Polymers | 2D $^{1}H$-$^{13}C$ INEPT | $B_0$=18.8 T; $\nu_{MAS}$=13 kHz; $\tau_{\pi/2 1H}$=10 μs; $\tau_{\pi/2 13C}$=8.3 μs; $\tau_J$=1.06 ms; $\nu_{dec}$=25 kHz; $\Delta t_1$=75 μs; $\tau_{t1}$=11.25 ms; $t_{1MODE}$=TPPI; $\tau_{acq}$=20.4 ms; $\tau_{dwell}$=10 μs; $n_{scan}$=16; $\tau_{recycle}$=1 s; T=16 ºC; | 1.3 h | $t_1$: 25 Hz/362 pts<br>$t_2$: 40 Hz/6144 pts |
| Uniform $^{13}C/^{15}N$ Tm1 Tail Domain-Only Polymers | 2D $^{15}N$-$^{13}C$ NCACX | $B_0$=17.5 T; $\nu_{MAS}$=12 kHz; $\tau_{\pi/2 1H}$=4 μs; $\tau_{\pi/2 13C}$=5 μs; $\nu_{CP1H}$=54 kHz; $\nu_{CP15N}$=32 kHz; $\tau_{CP}$=1.5 ms; $\tau_{\pi TPPM}$=5.6 μs; $\nu_{dec}$=83 kHz; $\nu_{SCP15N}$=17 kHz; $\nu_{SCP13C}$=29 kHz; $\nu_{13CCAR}$= 52.6 ppm; $\nu_{15NCAR}$=99.7 ppm; $\tau_{SCP}$=4.0 ms; $\tau_{DARR}$= 50 ms; $\nu_{DARR}$=12 kHz; $\Delta t_1$=82.6 μs; $\tau_{t1}$=5.62 ms; $t_{1MODE}$=States; $\tau_{acq}$=7.68 ms; $\tau_{dwell}$=15 μs; $n_{scan}$=1024; $\tau_{recycle}$=1.1 s; T=16 ºC; | 37.5 h | $t_1$: 0 Hz<br>$t_2$: 90 Hz |
| Uniform $^{13}C/^{15}N$ Tm1 Tail Domain-Only Polymers | 2D $^{15}N$-$^{13}C$ NCOCX | $B_0$=17.5 T; $\nu_{MAS}$=12 kHz; $\tau_{\pi/2 1H}$=4 μs; $\tau_{\pi/2 13C}$=5 μs; $\nu_{CP1H}$=54 kHz; $\nu_{CP15N}$=32 kHz; $\tau_{CP}$=1.5 ms; $\tau_{\pi TPPM}$=5.6 μs; $\nu_{dec}$=83 kHz; $\nu_{SCP15N}$=28.8 kHz; $\nu_{SCP13C}$=40.8 kHz; $\nu_{13CCAR}$=175.2 ppm; $\nu_{15NCAR}$=99.7 ppm; $\tau_{SCP}$=4.0 ms; $\tau_{DARR}$= 50 ms; $\nu_{DARR}$=12 kHz; $\Delta t_1$=82.6 μs; $\tau_{t1}$=5.62 ms; $t_{1MODE}$=States; $\tau_{acq}$=7.68 ms; $\tau_{dwell}$=15 μs; $n_{scan}$=1280; $\tau_{recycle}$=1.1 s; T=16 ºC; | 53.1 h | $t_1$: 0 Hz/188 pts<br>$t_2$: 90 Hz |

| | | | | |
|---|---|---|---|---|
| Uniform $^{13}C/^{15}N$ Tm1 Tail Domain-Only Polymers | 3D $^{15}N$-$^{13}C$ NCACX | $B_0$=17.5 T; $\nu_{MAS}$=12 kHz; $\tau_{\pi/21H}$=4 µs; $\tau_{\pi/213C}$=5 µs; $\nu_{CP1H}$=54 kHz; $\nu_{CP15N}$=32 kHz; $\tau_{CP}$=1.5 ms; $\tau_{\pi TPPM}$=5.6 µs; $\nu_{dec}$=83 kHz; $\nu_{SCP15N}$=16.8 kHz; $\nu_{SCP13C}$=28.8 kHz; $\nu_{13CCAR}$= 52.6 ppm; $\nu_{15NCAR}$=99.7 ppm; $\tau_{SCP}$=4.0 ms; $\tau_{DARR}$= 50 ms; $\nu_{DARR}$=12 kHz; $\Delta t_1$=106.2 µs; $\tau_{t1}$=5.63 ms; $t_{1MODE}$=States; $\Delta t_2$=129.8 µs; $\tau_{t2}$=4.54 ms; $t_{2MODE}$=States; $\tau_{acq}$=7.68 ms; $\tau_{dwell}$=15 µs; $n_{scan}$=32; $\tau_{recycle}$=1.1 s; T=16 ºC; | 72.6 h | $t_1$: 0 Hz<br>$t_2$: 20 Hz<br>$t_3$: 90 Hz |
| Uniform $^{13}C/^{15}N$ Tm1 Tail Domain-Only Polymers | 3D $^{15}N$-$^{13}C$ NCOCX | $B_0$=17.5 T; $\nu_{MAS}$=12 kHz; $\tau_{\pi/21H}$=4 µs; $\tau_{\pi/213C}$=5 µs; $\nu_{CP1H}$=54 kHz; $\nu_{CP15N}$=32 kHz; $\tau_{CP}$=1.5 ms; $\tau_{\pi TPPM}$=5.6 µs; $\nu_{dec}$=83 kHz; $\nu_{SCP15N}$=16.8 kHz; $\nu_{SCP13C}$=28.8 kHz; $\nu_{13CCAR}$=175.2 ppm; $\nu_{15NCAR}$=99.7 ppm; $\tau_{SCP}$=4.0 ms; $\tau_{DARR}$= 50 ms; $\nu_{DARR}$=12 kHz; $\Delta t_1$=106.2 µs; $\tau_{t1}$=4.2 ms; $t_{1MODE}$=States; $\Delta t_2$=129.8 µs; $\tau_{t2}$=3.9 ms; $t_{2MODE}$=States; $\tau_{acq}$=7.68 ms; $\tau_{dwell}$=15 µs; $n_{scan}$=64; $\tau_{recycle}$=1.1 s; T=16 ºC; | 93.9 h | $t_1$: 0 Hz<br>$t_2$: 20 Hz<br>$t_3$: 90 Hz |
| 50% [$^{13}C/^{14}N$] and 50% [$^{12}C/^{15}N$] Tm1 Tail Domain-Only Polymers | 2D $^{15}N$-$^{13}C$ zf-TEDOR | $B_0$=18.8 T; $\nu_{MAS}$=10 kHz; $\tau_{\pi/21H}$=3 µs; $\tau_{\pi/213C}$=4 µs; $\tau_{\pi/215N}$=6 µs; $\nu_{CP1H}$=62 kHz; $\nu_{CP13C}$=42 kHz; $\tau_{CP}$=1.0 ms; $\tau_{\pi SPINAL}$=6.3 µs; $\nu_{dec}$=83 kHz; $\tau_{gauss13C}$=350 µs; $\tau_{TEDOR}$=6 ms; $\tau_{z\text{-}filter}$=300 µs; $\nu_{z\text{-}filter}$=10 kHz; $\Delta t_1$=100 µs; $\tau_{t1}$=7.5 ms; $t_{1MODE}$=States-TPPI; $\tau_{acq}$=7.68 ms; $\tau_{dwell}$=5 µs; $n_{scan}$=2720; $\tau_{recycle}$=1 s; T=5 ºC; | 113.3 h | $t_1$: 100 Hz<br>$t_2$: 80 Hz |
| Segmental $^{13}C/^{15}N$ Tm1 Intermediate Filaments | 1D $^{13}C$ CP | $B_0$=18.8 T; $\nu_{MAS}$=13 kHz; $\tau_{\pi/21H}$=2.5 µs; $\nu_{CP1H}$=78 kHz; $\nu_{CP13C}$=63 kHz; $\tau_{CP}$ =1.0 ms; $\tau_{\pi SPINAL}$=5.3 µs; $\nu_{dec}$=100 kHz; $\tau_{acq}$=5.12 ms; $\tau_{dwell}$=5 µs; $n_{scan}$=1024; $\tau_{recycle}$=1 s; T={16, −2, −12, −19} ºC; | 0.3 h | $t_1$: 100 Hz |
| Segmental $^{13}C/^{15}N$ Tm1 Intermediate Filaments | 2D $^{13}C$-$^{13}C$ CP-DARR | $B_0$=18.8 T; $\nu_{MAS}$=13 kHz; $\tau_{\pi/21H}$=2.5 µs; $\tau_{\pi/213C}$=4 µs; $\nu_{CP1H}$=78 kHz; $\nu_{CP13C}$=62 kHz; $\tau_{CP}$ =1.0 ms; $\tau_{\pi TPPM}$=4.9 µs; $\nu_{dec}$=100 kHz; $\tau_{DARR}$= 50 ms; | 39.1 h | $t_1$: 160 Hz/312 pts<br>$t_2$: 160 Hz |

| | | | | |
|---|---|---|---|---|
| | | $\nu_{DARR}$=13 kHz; $\Delta t_1$=22.2 μs; $\tau_{t1}$=2.22 ms; $t_{1MODE}$=TPPI; $\tau_{acq}$=5.12 ms; $\tau_{dwell}$=5 μs; $n_{scan}$=352; $\tau_{recycle}$=2.0 s; T=−19 ºC; | | |
| Segmental $^{13}C/^{15}N$ Tm1 Intermediate Filaments | 1D $^{13}C$ INEPT | $B_0$=18.8 T; $\nu_{MAS}$=13 kHz; $\tau_{\pi/21H}$=10 μs; $\tau_{\pi/213C}$=4 μs; $\tau_J$=1.06 ms; $\nu_{dec}$=25 kHz; $\tau_{acq}$=10.2 ms; $\tau_{dwell}$=5 μs; $n_{scan}$=1024; $\tau_{recycle}$=1 s; T={16, −2, −12, −19} ºC; | 0.3 h | $t_1$: 50 Hz |
| Segmental $^{13}C/^{15}N$ Intermediate Filaments | 2D $^1H$-$^{13}C$ INEPT | $B_0$=18.8 T; $\nu_{MAS}$=13 kHz; $\tau_{\pi/21H}$=10 μs; $\tau_{\pi/213C}$=4 μs; $\tau_J$=0.994 ms; $\nu_{dec}$=25 kHz; $\Delta t_1$=75 μs; $\tau_{t1}$=11.25 ms; $t_{1MODE}$=States-TPPI; $\tau_{acq}$=30.72 ms; $\tau_{dwell}$=5 μs; $n_{scan}$=64; $\tau_{recycle}$=1 s; T=16 ºC; | 5.3 h | $t_1$: 25 Hz/212 pts<br>$t_2$: 40 Hz |
| Uniform $^{13}C/^{15}N$ Tm1 Tail Domain Ether Precipitated | 2D $^{13}C$-$^{13}C$ CP-DARR | $B_0$=17.5 T; $\nu_{MAS}$=12 kHz; $\tau_{\pi/21H}$=4 μs; $\tau_{\pi/213C}$=5 μs; $\nu_{CP1H}$=52 kHz; $\nu_{CP13C}$=38 kHz; $\tau_{CP}$ =1.5 ms; $\tau_{\pi TPPM}$=5.6 μs; $\nu_{dec}$=83 kHz; $\tau_{DARR}$= 50 ms; $\nu_{DARR}$=12 kHz; $\Delta t_1$=22.4 μs; $\tau_{t1}$=3.36 ms; $t_{1MODE}$=States; $\tau_{acq}$=7.68 ms; $\tau_{dwell}$=15 μs; $n_{scan}$=384; $\tau_{recycle}$=1.5 s; T=16 ºC; | 48 h | $t_1$: 90 Hz/212 pts<br>$t_2$: 90 Hz |

*$B_0$, magnetic field; $\nu_{MAS}$, sample spinning frequency; $\tau_{\pi/21H}$, $^1H$ π/2 pulse length; $\tau_{\pi/213C}$, $^{13}C$ π/2 pulse length; $\tau_{gauss13C}$, $^{13}C$ gaussian selective pulse length; $\tau_{\pi/215N}$, $^{15}N$ π/2 pulse length; $\nu_{CP1H}$, $^1H$ RF power during CP; $\nu_{CP13C}$, $^{13}C$ RF power during CP; $\nu_{CP15N}$, $^{15}N$ RF power during CP; $\tau_{CP}$, $^1H$-$^{13}C$ or $^1H$-$^{15}N$ CP contact time; $\tau_{z\text{-filter}}$, TEDOR z-filter time; $\tau_{TEDOR}$, TEDOR mixing time; $\nu_{z\text{-filter}}$, $^1H$ RF pulse during TEDOR z-filter; $\tau_{\pi TPPM}$, TPPM π pulse length; $\tau_{\pi SPINAL}$ SPINAL64 π pulse length; $\nu_{dec}$, $^1H$ decoupling RF power; $\nu_{SCP15N}$, $^{15}N$ RF power during SCP; $\nu_{SCP13C}$, $^{13}C$ RF power during SCP; $\tau_{SCP}$, $^{13}C$-$^{15}N$ SCP contact time; $\nu_{13CCAR}$, $^{13}C$ carrier frequency during SCP; $\nu_{15NCAR}$, $^{15}N$ carrier frequency during SCP; $\tau_{DARR}$, $^{13}C$-$^{13}C$ DARR mixing time; $\nu_{DARR}$, $^{13}C$-$^{13}C$ DARR RF power; $\tau_J$, J-delay for INEPT; $\Delta t_1$, $t_1$ increment; $\tau_{t1}$, total $t_1$ evolution time; $\Delta t_2$, $t_2$ increment; $\tau_{t2}$, total $t_2$ evolution time; $t_{1MODE}$, $t_1$ acquisition mode; $t_{2MODE}$, $t_2$ acquisition mode; $\tau_{acq}$, acquisition time; $\tau_{dwell}$, dwell time; $n_{scan}$, number of scans; $\tau_{recycle}$, recycle delay time; T, sample temperature;

**Frequencies corresponds to the amount of gaussian line broadening applied in the given spectral dimension. If indicated, 'pts' is the number of points used for linear prediction.
